# Specific Sensitivity to Rare and Extreme Events: Quasi-Complete Black Swan Avoidance vs Partial Jackpot Seeking in Rat Decision-Making

**DOI:** 10.1101/2021.11.01.466806

**Authors:** Mickaël Degoulet, Louis-Matis Willem, Christelle Baunez, Stéphane Luchini, Patrick A. Pintus

## Abstract

Most behavioral studies in animals investigate risk using outcome probabilities larger than 10%. However, real-world Decision-Making often requires evaluating events that are both extremely unlikely and highly consequential. To address this gap, we developed an experimental and computational framework to quantify how rats detect and adapt to **rare (probability < 1**%**) and extreme (deviation from mean > 10 standard deviations) outcomes (REE)**. Using a four-armed bandit task, animals chose between options associated with probabilistic rewards (sugar pellets) or punishments (time-out delays). Depending on the animal’s choice, REE can occur or not, allowing us to probe how rats integrate information across both common and fat-tailed event distributions. Across subjects, behavior showed **restricted diversification** (typically two out of four options) and clear **sensitivity to REE**, expressed as a systematic avoidance of rare and extreme punishments (“Black Swans”) combined with partial exposure to rare and extreme gains (“Jackpots”). The dominant behavioral phenotype displayed a near-complete suppression of exposure to Black Swans, whereas exposure to Jackpots remained only moderate. This asymmetric sensitivity came at a cost: these animals accepted smaller frequent gains and larger frequent losses to avoid catastrophic outcomes. To account for these behaviors, we implemented an **augmented reinforcement learning model** in which REE are weighted separately from frequent events. Fitting this model to individual behavioral data captured subjects’ decision patterns far better than standard Q-learning, which systematically failed to reproduce the observed asymmetry. The best-fitting model suggests that the rat brain segregates information from the central outcome distribution and the distribution tails, assigning distinct value weights to REE during action selection. This work provides the first evidence that rodents spontaneously adapt their learning strategies to avoid extreme punishments while partially maintaining exposure to rare and extreme gains, suggesting that frequent outcomes are treated separately from rare and extreme ones through different weights in Decision-Making.

## Introduction

Exploration and exploitation are fundamental components of adaptive behavior under uncertainty in both humans and non-human animals (for review see [28]). However, relatively little is known about how these behavioral processes are modulated by outcomes that are both extremely rare and highly consequential - hereafter referred to as *Rare and Extreme Events* (REE). Although risk processing has been extensively studied across psychology, behavioral economics, and neuroscience, most laboratory decision-making tasks rely on outcome distributions in which event probabilities exceed 10% (e.g. [9], [10], [41], [53]; see also [20], [27], [32] for notable exceptions). This represents a major limitation considering that in natural environments, many species must cope with events that occur far less frequently yet exert disproportionately large negative or positive impacts. Such events have shaped evolutionary trajectories and can even lead to species-wide extinction [19], while in humans they have occasionally driven technological and cultural leaps [29]. Because their probabilities and magnitudes cannot be reliably estimated from experience, REE pose profound challenges for learning and decision-making mechanisms. It remains largely unknown whether, and how, animals - including humans - detect, integrate, or adapt to such events.

Here, we address this gap by studying rats, a species widely used in behavioral and systems neuroscience to investigate the neural substrates of cost-benefit learning and decision-making (e.g. [6], [1], [49], [51], [24], [33], [47], [21], [58]; see [23] for relevance to human decision processes). We developed a novel four-armed bandit task in which rats repeatedly interact with their environment and encounter REE with probabilities *<* 1%. Critically, these REE produce outcome magnitudes - both gains and losses - that exceed 10 standard deviations relative to the average outcome. Standard rodent analogs of the Iowa Gambling Task typically use lower-magnitude rare events with probabilities ≥ 10% ([4], [1], [37], [57]). The central feature of our task is that it allows us to test whether rats adopt behavioral strategies that minimize exposure to extreme losses (Black Swans) while preserving the possibility of extreme gains (Jackpots) - analogous to “Anti-fragile” strategies described in conceptual work [48]. In contrast, strategies that eliminate Jackpots while maintaining exposure to Black Swans are here termed “Fragile”. Operationally, complete exposure to Jackpots and complete avoidance of Black Swans defines the Anti-fragile option, whereas the inverse pattern defines the Fragile option. We additionally define the Robust option, as the one excluding all REE and Vulnerable option the one exposing animals to both positive and negative REE. This framework is compatible with evolutionary hypotheses proposing that organisms may be biased toward choice policies that enhance long-term viability under uncertainty.

Our design separates outcomes into three probabilistic domains: (i) “normal events” (NE, ≈ 90% probability), (ii) “rare events” (RE, ≈ 10%), and (iii) REE (*<* 1%). Reward magnitudes scale nonlinearly with decreasing probability such that convex options produce increasingly larger gains as probability decreases (i.e., exposure to Jackpots) while concave options limit the possible gain within a short range whatever the probability (i.e. avoidance of Jackpots). In contrast, in the negative domain, concave options produce increasingly larger losses (exposure to Black Swans), while convex ones limit the possible losses within a short range whatever the probability. Convexity here refers not to utility curvature but to the accelerating relationship between magnitude and rarity.

Based on stochastic dominance arguments, value-maximizing agents with non-decreasing value functions should prefer concave options in the NE domain (first-order dominance), and continue to do so in the RE domain if they also display risk aversion (second-order dominance). However, in the full environment including the REE, second-order dominance reverses, and convex options become optimal because avoiding Black Swans is increasingly costly when REE fail to materialize. Thus, Anti-fragile strategies are only advantageous if REE are internally represented.

We incorporate both gains (sugar pellets) and losses (time-out punishment) to capture sensitivity across valence domains. We derive two key behavioral metrics: (1) *Total Sensitivity to REE*, defined as the combined frequency of convex choices in gain and loss domains, indexing whether rats incorporate REE into their strategy; and (2) *One-Sided Sensitivity to REE*, defined as the difference between convex gain choices and convex loss choices, quantifying in particular asymmetric Jackpot-seeking versus Black Swan avoidance.

Using 20 rats (about 6000 trials per subject over 41 sessions), we report two major behavioral findings. First, rats diversify their behavior across options but typically concentrate on roughly two of the four available options. Nineteen of 20 animals show moderate-to-high Total Sensitivity, indicating that most animals systematically combine some exposure to extreme rewards with partial avoidance of extreme losses. Second, 13 of 20 rats demonstrate near-complete avoidance of Black Swans while only partially seeking Jackpots, producing negative One-Sided Sensitivity and consistent choice mixtures of Anti-fragile and Robust options. This suggests differential weighting of rare positive versus rare negative outcomes. Consistent with this interpretation, both Total Sensitivity and Black Swan avoidance increase after rats directly experience a rare extreme loss. To interpret computationally these results, we evaluated a family of augmented Q-learning models that incorporate explicit sensitivity parameters for Jackpots and Black Swans. Model comparison using information criteria identifies an architecture in which NE+RE and REE are weighted separately during action selection, suggesting that rats assign distinct cognitive or neural salience to REE - especially negative ones - when making decisions under uncertainty.

## Results

### Measures of sensitivity to Rare and Extreme Events in a four-armed bandit task

Rats were tested in an operant chamber equipped with four nose-poke ports, each linked to a specific distribution of reward (sucrose pellets) and punishment (time-out delays) drawn from a fixed schedule across sessions (Fig. 1a). After training, animals completed 41 experimental sessions (20 min/session; ≈ 120 trials/session). As shown in Figure 1b, positive outcomes consisted of sucrose pellets, whereas negative outcomes corresponded to time-out periods during which ports were inactive, preventing pellet acquisition.

**Figure 1:**
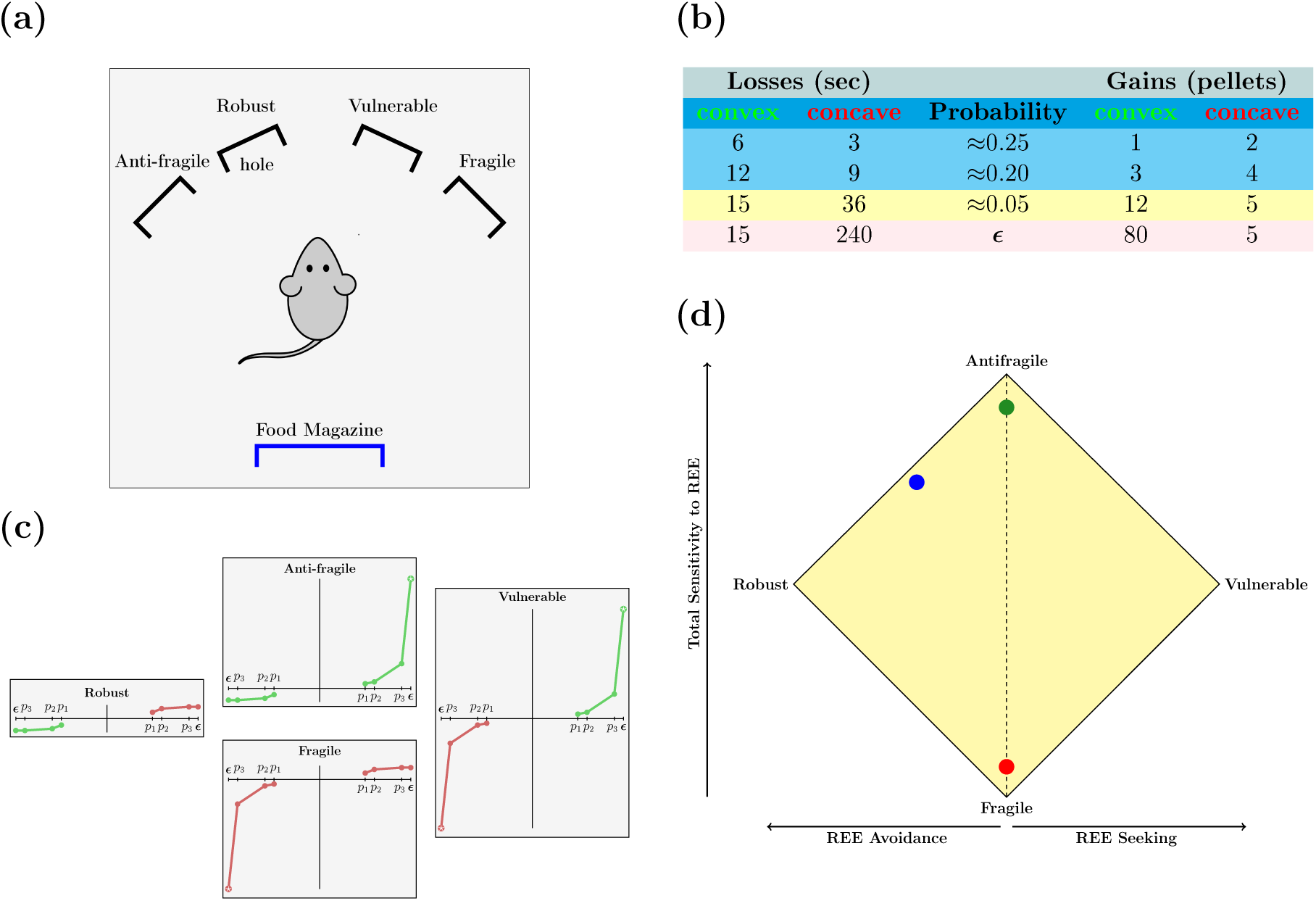
Experimental Design. (a) Schematic of the operant conditioning chamber. Rats performed a self-paced four-option choice task using nose-poke apertures, each corresponding to a distinct option leading to distribution of reward (sucrose pellets) or punishment (time-out delay) at various size and probability. (b) Outcome structure for each option. Rewards and losses follow either convex (green) or concave (red) probability-magnitude functions. Outcomes are grouped into three probability domains: high-probability Normal Events (NE; 20 − 25%, blue), Rare Events (RE; 5%, yellow), and Rare and Extreme Events (REE; *<* 1%, pink). (c) Functional characterization of the four choice options based on exposure to rare and extreme gains (“Jackpots”) and rare and extreme losses (“Black Swans”). On the horizontal x-axis are reported (in decreasing order moving away from the origin) the ex-ante probabilities that are unknown to the subject, which only observes the outcome that is measured on the vertical y-axis. For convenience, gains appear in the upper-right quadrant while losses appear in the lower-left one. Convex curve are in green while concave are red. Robust: convex losses and concave gains (avoids both REE). Anti-fragile: convex losses and convex gains (avoids Black Swans while retaining access to Jackpots). Fragile: concave losses and concave gains (exposure to Black Swans, avoidance of Jackpots). Vulnerable: concave losses and convex gains (exposed to both Black Swans and Jackpots). (d) Behavioral sensitivity metrics. Total Sensitivity (y-axis) quantifies overall incorporation of REE by summing convex choices in gain and loss domains. One-sided Sensitivity (x-axis) measures asymmetry between REE seeking and REE avoidance. Example profiles illustrate: an Anti-fragile-like pattern (green), a Fragile-like pattern (red), and a mixed Anti-fragile/Robust profile with strong Black Swan avoidance and partial Jackpot seeking (blue). See Material and Methods for computational definitions.

Each choice option was defined by a characteristic mapping between outcome magnitude and event probability. Outcome magnitudes could follow either convex (accelerating; green) or concave (decelerating; red) functions as event probability decreased (Fig. 1b-c). Critically, all options included a spectrum of outcome classes: highly probable Normal Events (NE; ≈ 90%), Rare Events (RE; ≈ 10%), and Rare and Extreme Events (REE; ≈ 1%), which corresponded to exceptionally large rewards (“Jackpots”: 80 pellets) or exceptionally large punishments (“Black Swans”: 240-s delay).

Combining convex or concave gain and loss functions generated four distinct options (Anti-fragile, Robust, Vulnerable, Fragile; Fig. 1c). The Anti-fragile option paired convex gains with convex losses, thereby allowing exposure to Jackpots while avoiding exposure to Black Swans. The Fragile option combined concave gains and concave losses, yielding small plateauing rewards but exposing animals to extreme losses (Black Swans). Robust and Vulnerable options showed complementary patterns: Robust avoided Black Swans but also prevented access to Jackpots, whereas Vulnerable exposed animals to both extreme outcomes.

Because the behavioral structure explicitly manipulated exposure to REE, we quantified how frequently each rat selected options containing convex (versus concave) outcome functions. Two measures were derived (Fig. 1d). Total Sensitivity to REE reflects the overall tendency to select options that allow contact with extreme outcomes (i.e., convex functions in either domain). One-sided Sensitivity quantifies asymmetry between the gain and loss domains - specifically, whether an animal preferentially seeks extreme rewards (Jackpot Seeking) or preferentially avoids extreme punishments (Black Swan Avoidance).

These two dimensions generate a behavioral space in which animals may cluster near extreme phenotypes (e.g., exclusively Anti-fragile or exclusively Fragile) or diversify among several strategies. Points lying between edges reflect mixed strategies. For example, a profile near the Anti-fragile - Fragile axis indicates strong Black Swan Avoidance combined with frequent-but not exclusive - Jackpot seeking.

Importantly, although Normal and Rare Events dominate the experience of the task, the inclusion of REE changes the adaptive landscape: extreme outcomes, despite their low probability, produce large shifts in reinforcement history. Thus, rats that behave as though REE “matter” (high Total Sensitivity) preferentially select options allowing contact with large rewards while minimizing catastrophic time-outs, whereas rats that behave as if REE are irrelevant (low Total Sensitivity) preferentially sample concave options that yield stable but limited outcomes, according to the stochastic dominance of order 1. Together, this framework allows quantification of how individual animals integrate extremely infrequent but high-impact outcomes into their decision strategy, enabling downstream analyses linking these behavioral phenotypes to neural processes.

### Main phenotype: strong avoidance of extreme punishments with partial sampling of extreme rewards

We analyzed the behavioral data collected across 41 daily sessions from 20 rats (see Material and Methods). To highlight the dominant features of the behavioral strategies adopted by the animals, Figure 2 plots each rat’s choices within the two-dimensional space defined by Total Sensitivity to REE and One-sided Sensitivity (relative weighting of extreme losses vs extreme rewards). This representation corresponds to the rotated square introduced in Figure 1d, where each vertex corresponds to exclusive selection of one of the four outcome distributions. A first inspection of Figure 2 shows that no animal operates at a single vertex - i.e., no rat commits exclusively to one contingency. Instead, all animals sample from multiple options, producing distributed behavioral profiles rather than categorical strategies. This diversification is qualitatively consistent with what is often observed in naturalistic foraging scenarios in which animals hedge against uncertainty or extreme events ([5], [40], [43], [48]).

**Figure 2:**
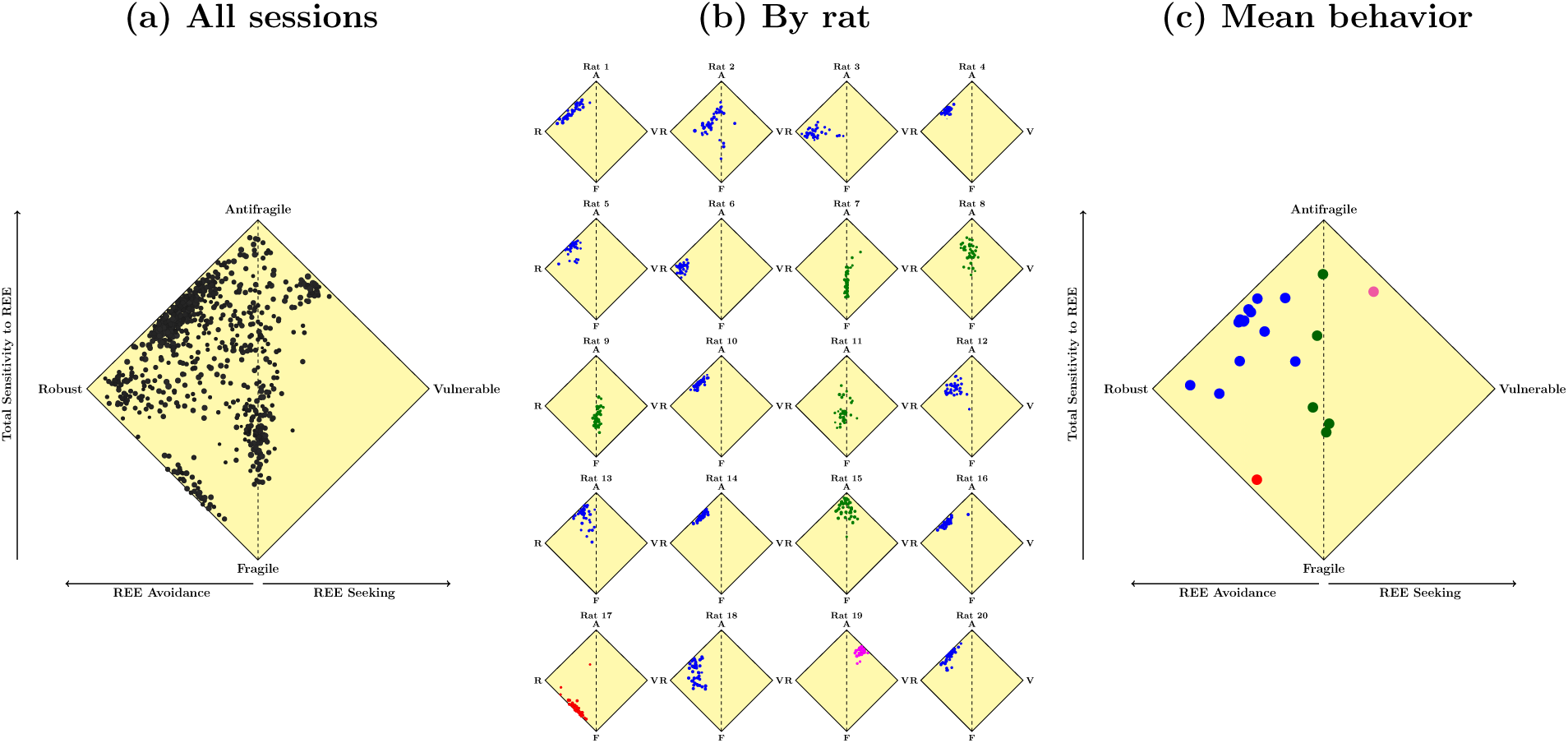
Total sensitivity to Rare and Extreme Events (REE) (y-axis) plotted against one-sided sensitivity to REE (x-axis), where rightward movement indicates REE seeking and leftward movement indicates REE avoidance. For a detailed description of the modeling and statistical analysis methods used, please refer to the Materials and Methods section. (a) the 820 black dots represent data from 41 sessions conducted with a total of 20 rats. (b) individual rat profiles, with each square corresponding to a single rat. The colored dots within each square denote the performance across the 41 sessions for each rat, categorized into two primary sensitivity profiles: blue dots indicate rats that exhibit strong sensitivity to REE and are classified as “Black Swan avoiders”, while green dots indicate rats that are responsive to REE but maintain a neutral stance, situated near the midline of the graph. Additionally, two outlier data points are highlighted in red and pink. (c) color-coded dots, each representing the average sensitivity measures across all 41 sessions per rat (n=20 rats).

### Population-level patterns

Panel 2a shows the full dataset (≈ 800 session-points). Most sessions lie on the left half of the behavioral space, corresponding to strategies that rather avoid REE. Within this domain, most sessions fall in the upper quadrant, indicating moderate to high Total Sensitivity - that is, rats systematically adjusted their choices in ways that reflect the presence of rare and extreme outcomes by minimizing exposure to extreme losses (Black Swans). Training history contributing to these patterns is detailed in Material and Methods.

### Individual behavioral profiles

Panel 2b displays each individual rat’s pattern across the 41 sessions. Each square corresponds to one rat, and the four vertices correspond to the four possible outcome distributions (A: Antifragile; R: Robust; V: Vulnerable; F: Fragile). With the exception of one animal (rat 17, red), all rats show moderate to high Total Sensitivity. Within this dominant group, two sub-groups emerge (blue and green). A second outlier, rat 19 (purple), is the only animal expressing both very high Total Sensitivity and strong Jackpot Seeking, achieved by alternating mainly between Antifragile and, less frequently, Vulnerable options. The remaining 18 animals fall into two stable phenotypes:

1. Blue group (13 rats): These animals primarily oscillate between the Robust and Antifragile options. This phenotype combines (i) moderate to high Total Sensitivity with (ii) pronounced avoidance of Black Swans. In other words, these rats preferentially avoid extreme punishments more than they seek extreme rewards.
2. Green group (5 rats): These rats primarily alternate between the Fragile and Antifragile options. They show moderate to high Total Sensitivity but display no directional bias in One-sided Sensitivity, indicating neither consistent avoidance of extreme losses nor active seeking of extreme rewards. Their choices are symmetric with respect to REE in the gain and loss domains.

### Average phenotypes

Figure 2c summarizes each rat’s average metrics across sessions. All but four rats exhibit an average Total Sensitivity above half of the maximum value; three are slightly below one, and one rat (rat 17) sits around 0.5, making it the closest to a Fragile-like pattern, though still deviating from true Fragile behavior. In total, 19 of 20 rats show clear sensitivity to the presence of REE.

Panel 2c also reveals a pronounced dominance of Black Swan Avoidance (negative Onesided Sensitivity) in the majority of animals. Rats 17 and 19 are opposite outliers: rat 17 has low Total Sensitivity, whereas rat 19 shows the highest Total Sensitivity coupled with strong Jackpot Seeking.

Within the 18-rat majority, the two main phenotypes described above are clearly distinguished: the green subgroup shows no One-sided bias, while the blue subgroup shows pronounced Black Swan Avoidance. Quantitative values are provided in Supplementary Material: Behavioral Measures per Rat.

### Structure in the relationship between Total and One-sided Sensitivity

A notable feature in panel 2c is a positive relationship between Total Sensitivity and One-sided Sensitivity among the 16 rats with the highest Total Sensitivity. Within this high-sensitivity group, animals that more strongly incorporate REE into their choices tend to show reduced Black Swan Avoidance (i.e., their One-sided Sensitivity becomes less negative). This shift reflects a progressive tilt toward convexity in the gain domain relative to the loss domain. Rat 19 represents an extreme case, approaching the maximal combination of Total Sensitivity and Jackpot Seeking.

This relationship is confirmed statistically: in the full dataset of 20 rats, the correlation between average Total and One-sided Sensitivity is near zero. However, restricting the analysis to the 16 high-sensitivity rats yields a correlation of ≈0.57. Over learning, this relationship strengthens: splitting the 41 sessions into four blocks of 10, the correlation increases from ≈0.17 initially to ≈0.56, ≈0.71, and ≈0.64 in successive epochs. Thus, the structure linking Total and One-sided Sensitivity emerges early and stabilizes over time.

### Behavioral measures of reward seeking and punishment avoidance

Figure 3 reports more intuitive behavioral metrics: the proportion of choices that (i) avoid Black Swans and (ii) seek Jackpots (detailed computation in Materials and Methods: Modeling and Statistical Analysis). Median values show a sharp dissociation between the two phenotypes. Approximately half of the blue rats avoid extreme punishments in *>* 90% of trials, whereas only ≈40% of green rats-and none of the outliers-achieve comparable avoidance.

**Figure 3:**
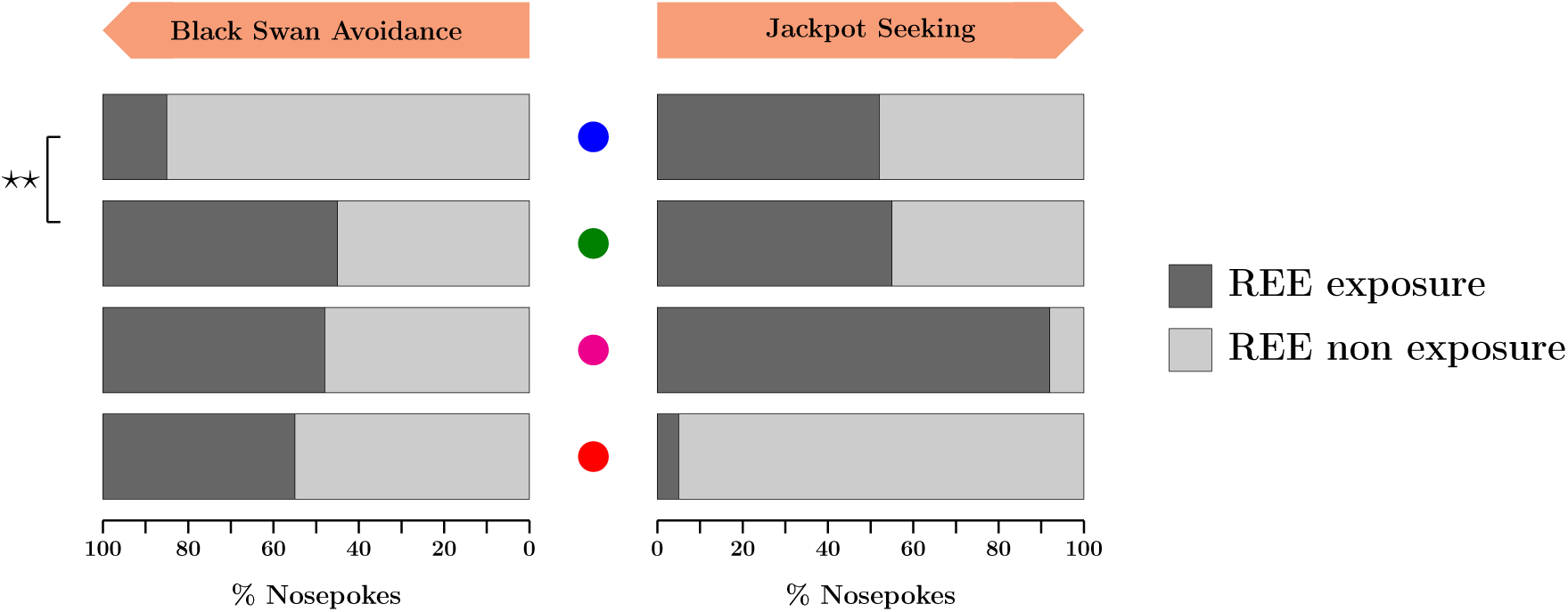
Behavioral Differences Between Phenotypes in Avoidance and Seeking Behaviors. This figure illustrates the median behavioral measures of Black Swan Avoidance and Jackpot Seeking across the two phenotypic groups, blue and green. The phenotypes are defined by distinct responses to Rare and Extreme Events (REE), with the blue phenotype exhibiting higher sensitivity to REEs and a tendency towards avoiding Black Swans, while the green phenotype shows less bias and remains closer to neutral behavior. Left Panel: This panel depicts the percentage of nosepokes where rats were exposed (black) or not exposed (light gray) to Black Swans. Increased Black Swan Avoidance is indicated by a higher proportion of non-exposure, suggesting a behavioral preference to avoid aversive outcomes among the blue phenotype compared to green. Right Panel: Conversely, this panel shows the percentage of nosepokes where rats were exposed (black) or not exposed (light gray) to Jackpots. Enhanced Jackpot Seeking is indicated by a higher proportion of exposure, suggesting greater motivation towards reward acquisition in the blue phenotype compared to green. Statistical comparisons between the phenotypes were conducted using Wilcoxon Rank Sum tests. Significant differences are denoted with *\*\** (*p <* 0.01).

By contrast, Jackpot Seeking does not differ between blue and green rats. Wilcoxon paired tests confirm this: blue rats differ significantly between punishment avoidance and reward seeking (p = 0.0002), whereas green rats do not (p = 0.6250). The two outliers diverge in opposite directions: rat 17 seeks almost maximal Jackpot exposure, while rat 19 nearly eliminates it. Exact values are given in the Supplementary Material.

### Black Swan outcomes strengthen sensitivity to rare and extreme events, whereas Jackpots do not

After establishing that most animals (19/20) display moderate-to-high Total Sensitivity to REE and robust avoidance of extreme punishments (Black Swans), we next asked how the actual experience of an REE shapes subsequent choice behavior within the same session. Specifically, we compared each rat’s behavioral sensitivity in the 10 nose-pokes preceding an REE with the 10 nose-pokes following it.

Figure 4 summarizes these within-session dynamics. For each rat (color-coded as in Figure 2), panel 4a shows the effects of experiencing a Jackpot, whereas panel 4b shows the effects of encountering a Black Swan. Total Sensitivity is plotted on the left column, and One-Sided Sensitivity (relative weighting of extreme losses vs gains) on the right. The dotted diagonal indicates no change.

**Figure 4:**
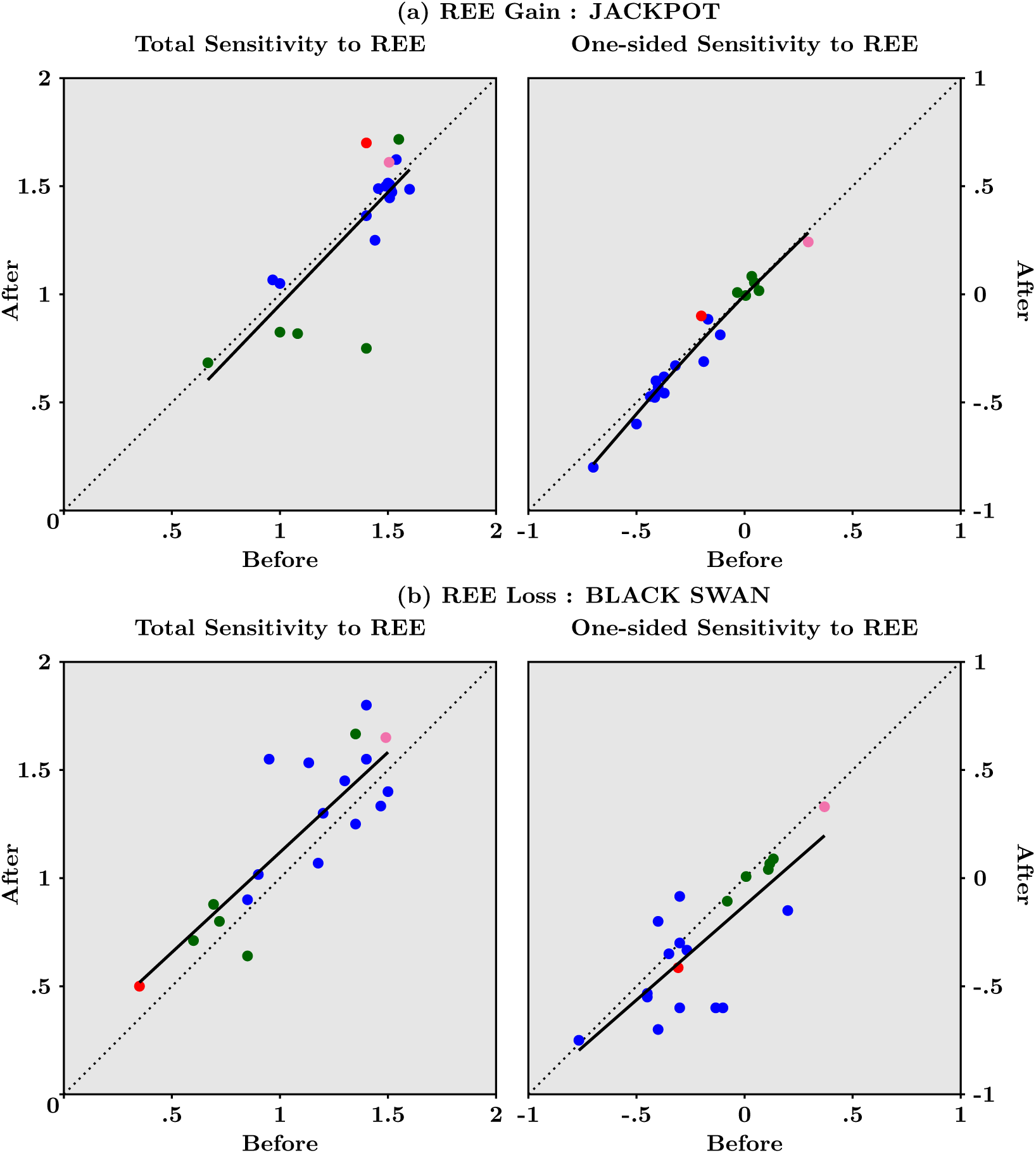
Responses of Total Sensitivity to REE and of One-sided Sensitivity to REE, averaged over the 41 final sessions, following Jackpots in panel (a) and Black Swans in panel (b), for each of the 20 rats. Each dot indicates one rat and dots are color coded to indicate the profile of the animal, as in Figure 2. Black dotted lines materialize no change in sensitivities after exposure to the REE compared to before. Black solid lines represent spline estimates - see Material and Methods.

A clear asymmetry emerges: Jackpots produce minimal change in either metric, with most points lying close to the diagonal. In contrast, Black Swan outcomes shift behavior systematically. Most animals show an increase in Total Sensitivity - i.e., they become more responsive to the structure of rare events-along with a decrease in One-Sided Sensitivity, reflecting a further shift toward avoidance of extreme losses. In both measures, points cluster above (Total Sensitivity) or below (One-Sided Sensitivity) the 45*^◦^* line, indicating a consistent strengthening of REE-related behavioral weighting after punishment but not after reward.

Thus, rare and severe losses - but not rare gains-acutely potentiate the animals’ sensitivity to REE. This pattern aligns with our overall finding that rats prioritize avoidance of extreme negative outcomes and that negative REE have disproportionate influence on choice updating. Bootstrapped statistics and additional tests reported in Material and Methods (see Table 3) confirm the significance of these effects.

**Table 1:**
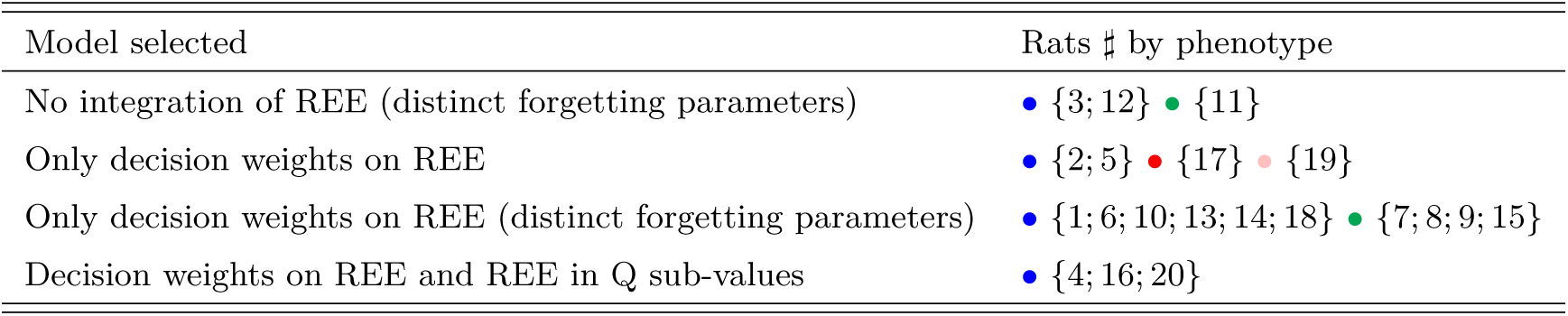
Selected augmented Q-Learning models by phenotype.

| Model selected | Rats # by phenotype |
| --- | --- |
| No integration of REE (distinct forgetting parameters) | • {3; 12} • {11} |
| Only decision weights on REE | • {2; 5} • {17} • {19} |
| Only decision weights on REE (distinct forgetting parameters) | • {1; 6; 10; 13; 14; 18} • {7; 8; 9; 15} |
| Decision weights on REE and REE in Q sub-values | • {4; 16; 20} |

**Table 2:** Statistical moments and decomposition of Jensen gaps for convex and concave exposures.

|  | Convex Exposure | Concave Exposure |
| --- | --- | --- |
| Mean | 3.89 | 3.11 |
| Std Deviation | 9.37 | 1.14 |
| Skewness | 7.16 | 0.20 |
| Kurtosis | 57.85 | 1.38 |
| $\log(\mathcal{JG})$ | 75.68 | 2.93 |
| $m_2$ (%) | $10^{-32}$ | 3.48 |
| $m_3$ (%) | $10^{-30}$ | 11.14 |
| $m_4$ (%) | $10^{-29}$ | 18.31 |
| $m_5$ (%) | $10^{-28}$ | 20.65 |
| $m_6$ (%) | $10^{-27}$ | 17.96 |
| $m_7$ (%) | $10^{-26}$ | 12.84 |
| $m_8$ (%) | $10^{-25}$ | 7.86 |
| $m_{80}$ (%) | 4.45 | $10^{-66}$ |

**Table 3:** Before/after differences for Total Sentivity to REE and for One-sided Sensitivity to REE, averaged over the 41 sessions, for each of the 20 rats - see Figure 4. We highlight negative mean differences in red and positive mean differences in yellow.

| Rat | Total Sensitivity to REE |  | One-sided Sensitivity to REE |  |
| --- | --- | --- | --- | --- |
|  | jackpot | black swan | jackpot | black swan |
| • 1 | -0.062 | -0.133 | -0.046 | -0.467 |
| • 2 | -0.025 | -0.108 | -0.075 | 0.215 |
| • 3 | 0.050 | 0.050 | -0.100 | -0.083 |
| • 4 | -0.036 | 0.100 | -0.036 | -0.300 |
| • 5 | 0.010 | 0.400 | 0.010 | -0.067 |
| • 6 | 0.100 | 0.117 | -0.100 | 0.017 |
| • 7 | -0.650 | 0.186 | 0.050 | 0.000 |
| • 8 | 0.017 | 0.111 | -0.050 | -0.044 |
| • 9 | -0.264 | -0.210 | 0.009 | -0.070 |
| • 10 | -0.045 | 0.400 | -0.009 | 0.200 |
| • 11 | -0.175 | 0.080 | 0.042 | -0.027 |
| • 12 | 0.033 | 0.150 | -0.122 | 0.000 |
| • 13 | 0.085 | 0.150 | 0.054 | -0.350 |
| • 14 | -0.114 | -0.100 | -0.086 | -0.500 |
| • 15 | 0.167 | 0.317 | -0.011 | -0.050 |
| • 16 | -0.000 | -0.100 | -0.062 | -0.100 |
| • 17 | 0.300 | 0.150 | 0.100 | -0.107 |
| • 18 | -0.190 | 0.600 | -0.010 | 0.000 |
| • 19 | 0.105 | 0.160 | -0.053 | -0.040 |
| • 20 | 0.014 | -0.100 | -0.029 | -0.300 |

In summary, within-session analyses reveal a strong valence asymmetry: Black Swan events transiently enhance both global sensitivity to REE and the bias to avoid rare losses, whereas Jackpots do not measurably alter behavioral sensitivity. This suggests that rare aversive outcomes exert a privileged influence on decision-making processes in this task, reinforcing the dominant phenotype of heightened vigilance toward extreme negative events.

### Q-Learning models require specific REE decision weights to mimic rats’ behavior

We have introduced sensitivity to Black Swans and Jackpots in a new class of augmented Reinforcement Learning models and we have estimated their parameters using observed choices and outcomes for each rat. The selected model ended up with a distinctive feature: it separates normal from rare and extreme outcomes through different weights in the Decision-Making process. Adding such specific sensitivity results in a good fit of the selected model - and simulated behaviors that are close - to behavioral observations, whereas a standard Q-learning model without sensitivity to REE is rejected for almost all rats.

To each of the four available options, indexed below by “*o*” (from 1 to 4, referring to Antifragile, Fragile, Robust, Vulnerable) in equations (1), we attach at each moment in trial/time *t* a gain sub-value *Q^g^*(*o*) and a loss sub-value *Q^l^*(*o*) that are updated as follows:

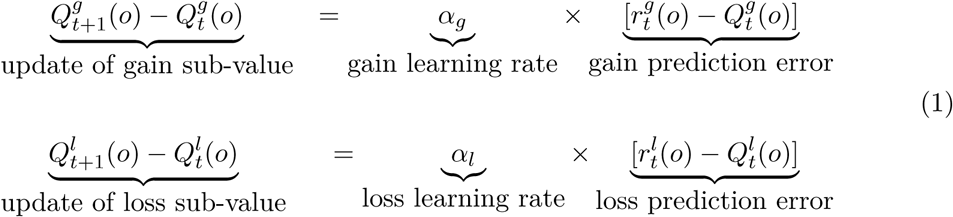

where the convention that updated values are indexed by *t* + 1 means right after observing the gain r_t_^g^ or loss r_t_^l^ corresponding to the choice made at *t*. Parameters *α_g_* and *α_l_* are usually labeled learning rates, indicating the speed at which reward prediction error gets updated into the latest values. Limiting cases occur when setting the learning rate either to zero - its smallest possible value - which means no subvalue updating, or to one - its largest possible value - which means that the subvalue tracks the obtained reward itself. While equations (1) applies for the updating of the chosen option, a similar updating rule applies for the three other options that are not chosen at that moment, by assuming that this happens as if the reward was zero, with different parameters. That is, for options that are not chosen, the updating rules for subvalues are simply 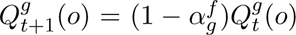 and 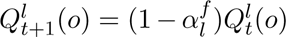, where the forgetting rates *α^f^_g_* and *α^f^_l_* are assumed to be bounded between zero and one.

The novel feature of our augmented Q-Learning model, besides integrating gains and losses, is that specific weights are attached to options that may produce REE. More specifically, we model the value of each option *o* as:

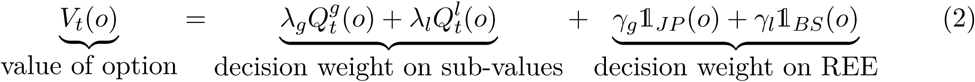

In equation (2), the value of each option *V_o_* is the sum of two terms. The first is the decision weight attached that sums up the gain and loss subvalues, each weighted by parameters *λ_g_*and *λ_l_* that may reflect a differential effect. Importantly, in the class of nested models that we estimated, REE may or may not be incorporated in the gain and loss subvalues, that should be thought of as averages - though not arithmetic but of an exponential moving type in view of equations (1) - see page 32 in [50]. The second and key term in equation (2) specifically captures the decision weight attached to REE, and it can itself be decomposed into the decision weights on Jackpots and on Black Swans. More precisely, the indicator function 1*_JP_* (*o*) equals one if the option exposes to Jackpots (namely when the chosen option is either A or V) and zero otherwise. Similarly, 1*_BS_*(*o*) equals one if the option exposes to Black Swans (namely options F and V) and zero otherwise. Therefore, *γ_g_* and *γ_l_* reflect the (possibly different) weights attached to Jackpots and Black Swans in the decision to choose one particular option rather than any other available.

Finally, the associated so-called action probabilities are given, for each option, by the Softmax function (a.k.a. the multinomial Logit model) 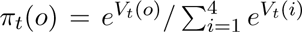, each a number between zero and one.

In our model-fitting procedure (described in detail below), the key parameters of interest are the decision-weight terms for rare and extreme gains (*γ_g_*) and losses (*γ_l_*). In the baseline Q-learning model, both terms are fixed at zero, meaning that REE do not receive any special weighting beyond their contribution to standard value updates. We therefore compared two families of candidate models to this benchmark: one in which REE contribute directly to option sub-values, and another in which REE influence choice only through dedicated decision-weight parameters (*γ_g_*,*γ_l_*).

The latter class of models is especially informative. If selected, it suggests that animals treat REE as qualitatively distinct inputs to the decision process-processed separately from average reward estimates-rather than integrating them into conventional value updating. Overall, the augmented Q-learning framework includes up to eight free parameters: learning rates for gains and losses, forgetting rates for gains and losses, decision weights for gain- and loss-related sub-values, and the two REE-specific decision weights *γ_g_* and *γ_l_*.

Model comparison using information criteria (summarized by phenotype in Table 1) shows that 17 of 20 animals incorporate REE using dedicated decision weights, whereas only three animals are best fit by models in which REE carry no special computational status. Most rats (including 8 blue, 4 green, and the red and pink animals) rely specifically on *γ_g_* and/or *γ_l_* to shape choices, with or without distinct forgetting parameters. A minority of blue rats incorporate REE both through decision-weight parameters and through REE contributions to sub-values. Learning and forgetting rates did not systematically distinguish between model classes.

Figure 5 presents the predicted sensitivities (bottom left) and simulated sensitivities (bottom right), compared with empirically observed sensitivities (top; identical to Figure 2, right panel). Predicted sensitivities - computed using each rat’s estimated parameters and the actual reward sequence - closely reproduce empirical sensitivity patterns. Simulated sensitivities were obtained by running each selected model through artificial 41-session datasets, using median parameters for the blue and green phenotypes (and individual parameters for the red and pink rats). These simulations show that the selected models generate stable behavioral phenotypes for the blue, red, and pink animals. Green rats, however, tend to segregate into two stable subgroups along the vertical (One-Sided Sensitivity) axis.

**Figure 5:**
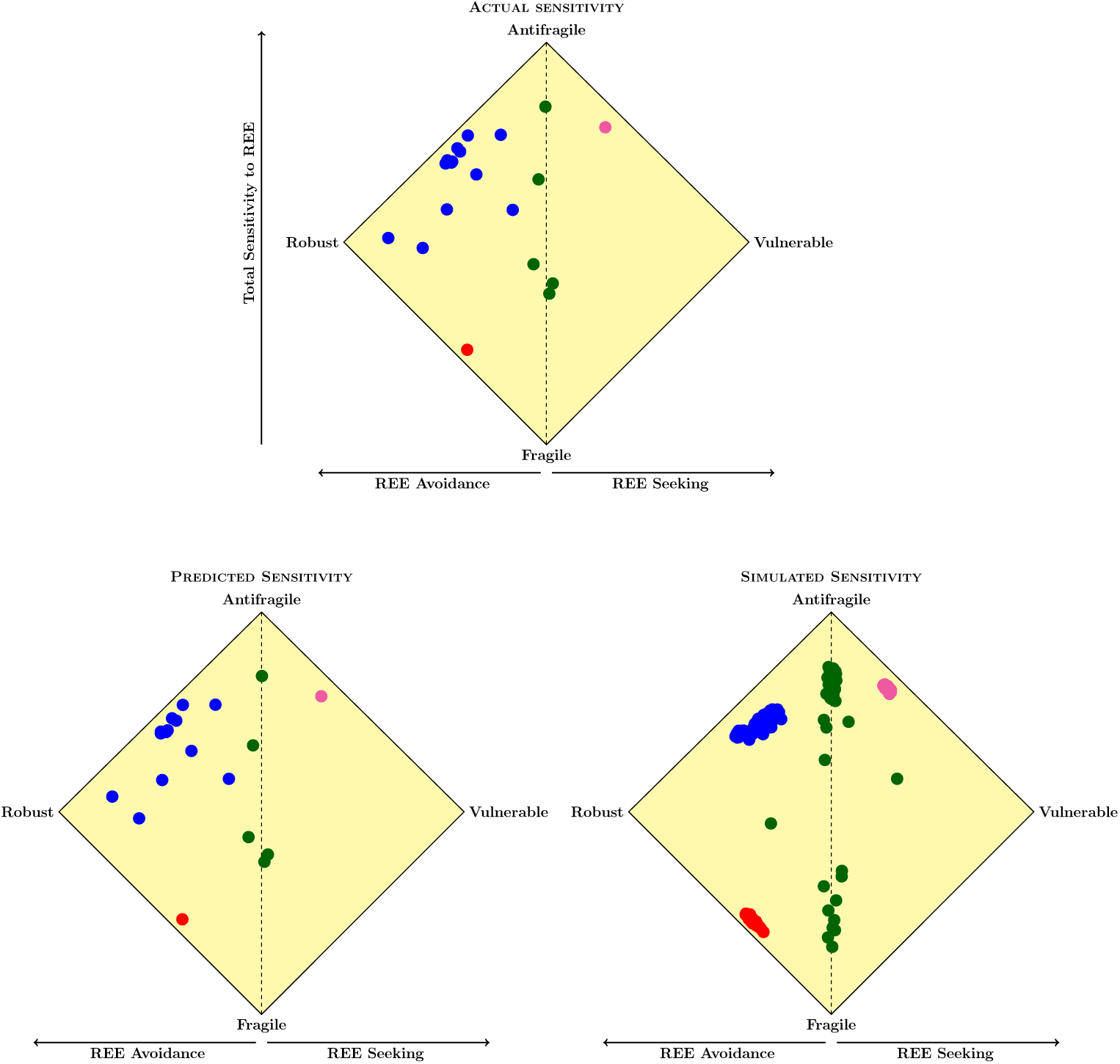
**Total Sensitivity to REE (*y*-axis) against One-sided Sensitivity to REE (*x*-axis)** - averages for each rat over the 41 sessions; top panel replicates the right panel in Figure 2: observed sensitivities; bottom left panel: predicted sensitivities derived from selected models; bottom right panel: simulated sensitivities computed from runs of selected models over artificial 41 sessions

Estimated parameters for each rat are shown in Figure 6. A prominent distinction between phenotypes emerges when focusing on the REE-specific decision weights: blue rats show a strong selective sensitivity to Black Swan, expressed as significantly negative *γ_l_* values (p = 0.0102). This indicates a robust computational penalty assigned to options that expose the animal to Black Swans (options F and V), consistent with their observed behavioral avoidance of these outcomes. In contrast, green rats do not show this selective aversive weighting and typically combine options F and A, suggesting a less selective treatment of REE. The before/after analysis illustrated in Figure 4 shows that the behavioral response to Black Swans is locally small in terms of both Total and One-sided sensitivities. This suggests that such effects are likely to be too subtle to be captured by this class of models for most rats.

**Figure 6:**
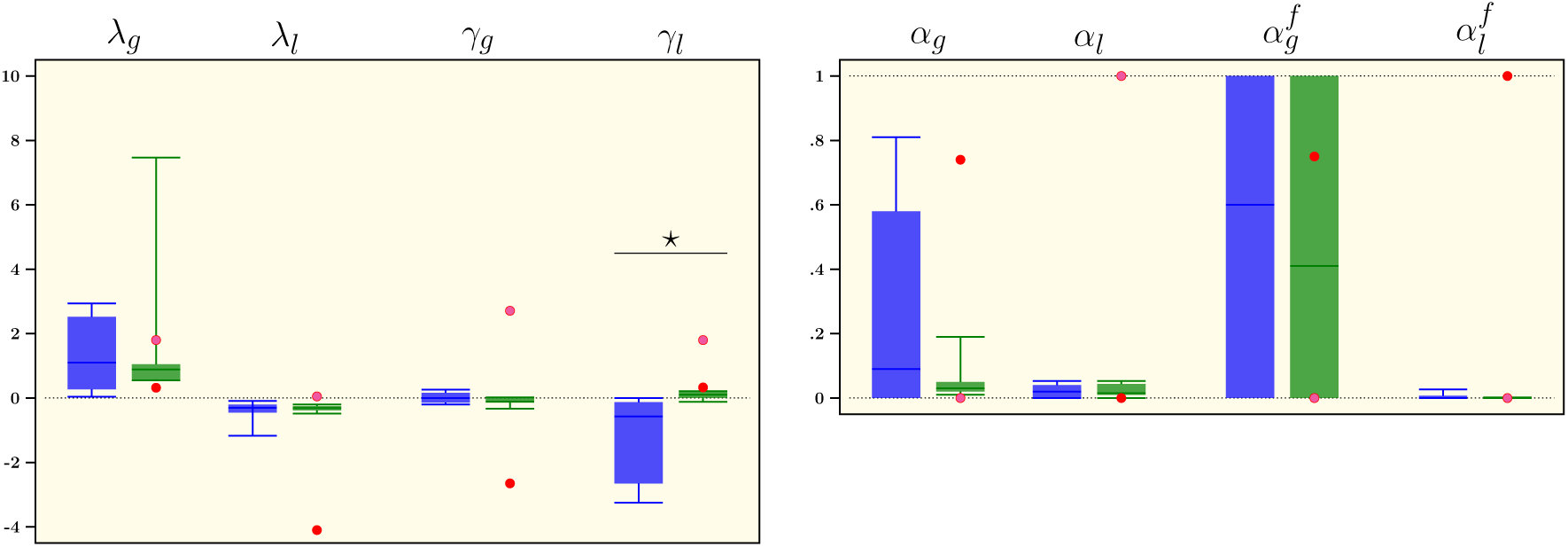
Parameters of the selected Q-Learning model, estimated for each rat over the 41 sessions - left panel: decision weights for averages of gains and losses and for REE; right panel: learning and forgetting rates for gains and losses; while individual parameter for pink and red rats are represented as dots, parameters for blue and green rats are represented by box plots with 10th and 90th percentiles, interquartile ranges and median; between phenotype comparisons carried out by Wilcoxon Rank Sum tests (*\** means p < 0.05) - see Material and Methods subsection Augmented Q-Learning Model Estimation and Simulation for details

## Discussion

### Asymmetric behavioral responses to rare extreme losses and rare extreme gains

We showed, first, that all rats in our sample exhibit some degree of Total Sensitivity to REE, with most animals displaying medium to high sensitivity. Second, we identified a dominant behavioral phenotype characterized by near-complete Black Swan Avoidance combined with partial Jackpot Seeking. We interpret these findings as reflecting a fundamental asymmetry between negative and positive REE, with a few caveats. Though we cannot fully exclude the possibility that satiety contributed to the lack of complete Jackpot Seeking, we did not systematically observe animals that stopped seeking the Jackpot. Because we restricted the rare outcome to occur on either the 10th or the 60th activation in a session (Table S1 in Supplementary Material), the question of whether the animals learn this association may arise. If the animals had learnt the 10th and 60th activation, they would exhibit a choice strategy that would tend to be more optimized than what is observed. For example, the options offering the possibility to obtain the Jackpot are not optimal in terms of gains for the frequent events, therefore the animals should tend to select these options only around the 10th and 60th choice. Most of their other choices should favor the options delivering the larger gains in the frequent domain. This is not what is observed.

In our task, avoiding Black Swans requires accepting longer frequent delays, and seeking Jackpots requires foregoing larger frequent gains. However, rats can guarantee the avoidance of Black Swans by selecting convex loss schedules, making these options relatively attractive. In contrast, no sequence of choices can guarantee the occurrence of a Jackpot-its realization depends on rare stochastic draws. For illustration, if 150 choices are made in a session (within the observed range) and the ex-ante REE probability is 1%, a Poisson approximation gives a probability of ≈ 78% for observing exactly one REE-a non-negligible chance for zero occurrence. Thus, our design captures a general asymmetry likely to characterize positive versus negative REE more broadly: while extreme losses can in principle be fully avoided, extreme gains cannot be assured, even under persistent optimal seeking. This asymmetry has important consequences for choice behavior. Rats experience a continuous stream of evidence-derived from the frequent-outcome domain-that points toward concave options as locally optimal. In particular, the Fragile option yields higher frequent gains and smaller frequent losses, and stochastically dominates the other options when REE do not occur. Although Fragile is dominated by Anti-fragile when both the Black Swan and Jackpot occur, complete Black Swan Avoidance provides near-certain protection against extreme losses. This makes strategies combining Anti-fragile and Robust options attractive, despite violating Stochastic Dominance in the frequent domain: full-domain Stochastic Dominance is effectively secured.

In contrast, complete Jackpot Seeking is less attractive because it cannot ensure the occurrence of a Jackpot; the cost of receiving smaller gains on most trials becomes unjustified unless the compensatory extreme reward actually occurs. Partial Jackpot Seeking therefore emerges as a more balanced strategy: it reduces violations of Stochastic Dominance in the frequent domain while still allowing some exposure to large rare gains. Put simply, bearing frequent relatively large losses is acceptable only if the avoided Black Swan is certain, and bearing frequent small gains is acceptable only if the compensating Jackpot is likely to appear-which it is not. Our results indicate that the “blue” rats have learned this property, combining Robust and Anti-fragile options to achieve near-complete Black Swan Avoidance with partial Jackpot Seeking. In contrast, “green” rats continue to mix Anti-fragile and Fragile options, leaving them exposed to Black Swans.

Interestingly, since rats do not stick to one choice but show some flexibility in their strategy, this rules out the influence of a spatial bias as the main driver in their response. Interestingly indeed, the design of the task was biased in two ways: the favorite hole during training was allocated to the antifragile option as a proof of concept and the REE occurred at the 10th and 60th choice if the appropriate option was chosen. Since most rats diversified their choices, the biased location does not seem to have prevented rats to learn all options. One might wonder whether or not they may have understood and learnt the 10th and 60th choice REE delivery. If this was the case, a trend towards optimal choice such as options delivering the largest rewards and the lowest losses in the most frequent domain with a shift towards options with positive REE and avoidance of negative REE around the 10th and 60th trial should have been observed. This was never the case.

In theory, because frequent outcomes occur often, animals should be able to detect Stochastic Dominance in the frequent domain relatively quickly. However, prior work indicates that animals require many sessions to learn Stochastic Dominance (e.g. [59], [3]). Our findings suggest a possible explanation: substantial experience may be needed to infer that REE have not occurred and can be excluded from the estimated distribution.

At this stage, it is natural to ask whether our findings reflect loss aversion or ambiguity aversion. Although most studies do not focus on REE, tasks involving both positive and negative outcomes generally report loss aversion (e.g., [18], [14], [61]), consistent with Prospect Theory ([22]). Models based on this framework, however, tend to fit rat behavior more effectively in tasks involving only rewards, where losses correspond to the absence of a positive outcome rather than punishment ([10],[3]). Emotional and anxiety-related mechanisms may also contribute ([60]), potentially implicating the basolateral amygdala, which has been linked to differential evaluation of negative outcomes in rats ([54]). Ambiguity aversion in non-human animals remains relatively unexplored (but see [44]).

Our results tentatively suggest that rats may exhibit ambiguity reduction with respect to REE: combining near-complete Black Swan Avoidance with only partial Jackpot Seeking collapses the sampling probability of Black Swans toward zero, while keeping the sampling probability of Jackpots bounded between zero and *ε*. This behavioral asymmetry provides further evidence that extreme gains and losses receive distinct treatment during decision making. Classical Q-learning models, which allow differential weighting of gains and losses, fail to capture these patterns. In contrast, the augmented Q-learning models incorporating explicit sensitivity to REE accurately reproduce the observed behaviors – principally through the strong aversion to Black Swans that distinguishes the “blue” phenotype from the “green” one.

While a wavy recency effect has been documented in human experiments ([38]), the before/after analysis reported in Figure 4 suggests that there is no sizeable immediate recency effect for Jackpots in rats. Even for Black Swans, the immediate recency effect we report remains modest when using a 10-trial window, and the analysis of the choice immediately following a REE does not show evidence of immediate negative recency. Although further investigation is beyond the scope of this paper, possibly interacting recency effects of Jackpots and Black Swans seems a promising research avenue. Also note that the task design does not allow rats to completely avoid negative rare events (RE) unless they cease performing the task altogether - a pattern typically seen in paradigms involving aversive stimuli such as electric foot shocks. The fact that all 20 rats maintained stable performance across the 41 sessions therefore provides evidence against a pronounced “hot-stove effect” (see [12]).

### The augmented Q-learning models suggest a specific neural pathway for REE encoding

The close fit and predictive accuracy of the augmented Q-learning model for the dominant behavioral phenotype point to a novel neurobiological hypothesis. Within this modeling framework, 11 of the 13 rats in the blue group (as defined in the Materials and Methods) appear to exclude REE from the Q-value update, yet display a specific sensitivity to Black Swans through a negative parameter *γ_l_*. In other words, when evaluating available actions, ”blue” rats do not integrate Black Swans into the running average of losses; instead, they assign them a distinct decision weight, effectively reducing the subjective value of options that expose the animal to these extreme losses.

In humans, rare and extreme stimuli are known to be disproportionately salient in perception and memory compared to moderate ones (e.g., [26]). Our behavioral and modeling results suggest that rats may exhibit a similar cognitive bias. This, in turn, raises the possibility that REE - particularly rare and extreme losses-may rely on neural encoding mechanisms that differ from those that process frequent, small rewards, which are classically attributed to dopamine neurons ([9], [46], [61], [55]), the striatum ([7], [11]), and cortical regions such as the anterior cingulate and orbitofrontal cortex ([45], [15]).

Identifying where and how REE are encoded in the brain remains an open question. If integrating REE requires representing long-term or low-frequency outcomes-beyond the short-term reward horizons typically studied (e.g., [56]) - candidate regions may include the anterior insula and the amygdala. The latter, given its central role in emotional salience, is a plausible substrate for representing highly aversive and unpredictable outcomes, which are known to bias choice asymmetrically toward negative events ([7], [54]).

From a computational standpoint, it is statistically rational to avoid folding extreme outliers into the mean of a distribution, as this can destabilize the estimate. Analogously, adaptive decision-making may benefit from treating REE separately from frequent outcomes, and from imposing different weights on Black Swans versus Jackpots. If such differential encoding confers evolutionary advantages in volatile environments (see [8]), then we might expect dedicated neural mechanisms to have evolved-possibly across multiple species.

To our knowledge, this specific hypothesis, grounded jointly in behavioral data and computational modeling, has not yet been directly tested in neuroscience. Doing so would align with the computational-validity framework advocated in [42], and represents an important direction for future research.

### Relevance for humans

Whether humans would also find Black Swan Avoidance and Jackpot Seeking attractive - and how they might combine these strategies in a similar experimental context-is an obvious question raised by our results. Our experimental design can be readily adapted to human participants, allowing us to test whether people likewise tend to avoid harmful REE and pursue beneficial ones. We have started exploring modified designs for human subjects in which the rare but non-extreme outcomes are removed, in line with [27] who emphasize that combining rarity and extremity is key. Preliminary results indicate that the behavioral phenotypes observed in rats also emerge in humans under these modified conditions, suggesting that REE are the primary drivers of observed behaviors. Furthermore, such an extended task may be a promising tool to delineate decision-making profiles in professional environments (e.g. firemen) and also to investigate vulnerability to addiction in patients. This translational potential makes the present findings in rats relevant to broader issues concerning the past and future of humanity.

Within the framework of the Anthropocene, mounting evidence shows that climate change and other environmental disruptions driven by human activities generate extreme events (see e.g., [13]), including rare but potentially catastrophic ones that could threaten many species with partial or complete extinction. Although uncertainty remains about the magnitude of these risks, empirical data place humanity in a situation that raises difficult but pressing questions. Why has our species historically failed to avoid behaviors and policies that increase exposure to destructive REE? And to what extent will humans be able to cope with such events in the near future, should they occur? Put differently, are humans following trajectories that diverge from Black Swan Avoidance rather than moving toward it?

While it is intuitive to assume that avoidance of destructive REE-and the ability to cope with them when they occur - provides adaptive value, compelling evidence for such evolutionary mechanisms remains scarce (but see [8]). If correct, this idea introduces a paradox: humans disproportionately contribute to creating environments rich in harmful REE. Does our species possess a particular trait that impairs our ability to protect ourselves against them, despite the apparent evolutionary benefits of doing so? Because destructive REE are increasingly endogenously generated by human activity, one may even wonder whether a uniquely human developmental or epigenetic factor predisposes us to decisions that fail to eliminate exposure to extremely harmful outcomes. If so, such a factor could also impair our ability to cope with REE once they become unavoidable. For instance, humans might struggle to detect predictive cues of REE - such as convexity in our experimental design (see [2]).

The results from our rat experiments fortunately do not support this pessimistic view. The substantial individual variability we observe-where some animals fail to avoid rare but extreme losses even when given the opportunity - suggests that humans might share this variability rather than being uniquely deficient. Importantly, the majority of rats consistently chose options that nearly eliminated exposure to harmful REE while still allowing partial access to beneficial REE in the form of Jackpots. This “adaptive” subgroup constitutes the largest portion of our sample.

A speculative hypothesis follows: some humans, and perhaps many, may also adopt strategies that avoid harmful - and sometimes pursue beneficial - rare extreme outcomes. More broadly, if sensitivity and behavioral responses to REE vary across individuals within a species, this raises important questions about how such heterogeneity shapes population dynamics, social structures, and, crucially, collective decision-making.

## Material and Methods

### Experimental Model and Subject Details

#### Animal Subjects

Adult Lister Hooded males (n = 20, ≈200 g at arrival, Charles River) were housed in groups of two in Plexiglas cages and maintained on an inverted 12 h light/dark cycle (light onset at 7 pm) with water available *ad libitum*, in a temperature - and humidity - controlled environment. Food was slightly restricted (≈80% of daily intake). Animal care and use conformed to the French regulation (Decree 2010-118) and were approved by local ethic committee and the French Ministry of Agriculture under #03129.01.

### Experimental Method Details

#### Apparatus

All behavioral experiments took place during the animals’ dark phase in standard five-hole operant boxes (MedAssociates) located in ventilated sound-attenuating cubicles. One side of each box was equipped with a central house light, a tone generator and a food magazine, outfitted an infrared beam for detecting nose poke inputs. Sucrose pellets (20 mg; Bio-Concept Scientific) were delivered from an external food pellet dispenser. An array of five response holes was located on the opposite curved wall, each equipped with stimulus lights and infrared beams for detecting input (nose poke). The center hole was continuously closed throughout the experiments (Fig. 1a). Data were acquired on a PC running MedPC-IV.

#### Design of Behavioral Sequences

Four menus were elaborated by mixing convex and concave exposures for both the gains (sugar pellets) and the losses (time-out punishment) described on Figure 1 (panel (*b*)). For the gains, animals could obtain 1, 3 (NE domain, blue), 12 (RE domain, yellow) or 80 (REE domain; pink) sugar pellets in the convex exposure or 2, 4, 5 or 5 pellets in the concave exposure. For the losses, convex exposure may impose 6, 12, 15 or 15 sec of time-out punishment while it was 3, 9 (NE domain; blue), 36 (RE domain, yellow) or 240 sec (REE; pink) for the concave exposure.

The four behavioral options depicted in Figure 1, panel (*c*), are therefore combinations of the above convex and concave exposures: the “Anti-fragile” exposure at the top middle is convex in both gains and losses, while the “Robust” option (left) is only convex for the losses. On the other hand, the “Vulnerable” exposure (right) is convex only for the gains, while the “Fragile” option (bottom middle) is concave for both gains and losses. This implies that the extreme - but rare - gain of 80 pellets (i.e. Jackpot) may be delivered only when either the Anti-fragile or the Vulnerable options are picked, while the extreme and rare time-out punishment of 240 sec (i.e. Black Swan) may be only experienced if choosing either the Fragile or the Vulnerable options.

The first three events of each behavioral options belong to the frequent domain as their frequency of occurrence, respectively 0.5, 0.4 or 0.1, is significantly larger than zero. During behavioral testing, animals equivalently experienced the gain and loss domains. On the other hand, extreme outcomes, i.e. Jackpot or Black Swan, have a much smaller likelihood of occurring since they may appear only at particular point during the behavioral sequences (see below). This means that they could happen if a rat has chosen an exposure that is either convex in the gain domain or concave in the loss domain at a given time. This implies that the frequency of rare and extreme events is less than half of one percent for all rats. Despite their low probability, extreme events have some importance because of their value. Our calibration of both frequencies and outcomes ensures that concave exposures dominate convex exposures: if the extreme gain (Jackpot) does not materialize, the expected payoff in sugar pellets is larger for concave exposures than convex exposures (i.e. ex-post first order stochastic dominance, over the frequent domain). Similarly, the expected time-out punishment is lower for concave exposures compared to convex ones, if the Black Swan does not happen. However, ex-post first-order stochastic dominance is reversed in the presence of extreme events, in which case convex exposures become more interesting in terms of payoff than concave ones. The reason we imposed such a dominance reversal was as follows. If rats always choose concave exposures, then they show no sensitivity to rare and extreme events, since they act as if those events never occur and always go for first-order stochastic dominance. This gives us our first measure, Total Sensitivity to REE, that simply sums up the proportion of convex exposures that are chosen for each rat over the 41 sessions. Because we deliberately integrate gains and losses in our design, we need to distinguish whether rats tend to choose convex exposures symmetrically over the gain and loss domains. We say that a particular rat exhibits Black Swan avoidance when it picks convex exposures in the loss domain more often than in the gain domain. Likewise, Jackpot Seeking occurs when convex choices are more frequent in the gain domain.

To avoid potential learning of the event occurrence during behavioral training and testing, ten different sequences of events, with respect of the first-order stochastic dominance as well as the balance between gain and loss domain exposures, were generated and randomly used for behavioral training and testing (Supplemental Table S3). Furthermore, to increase the rarity and the unpredictable nature of extreme events, the sequences of events used during behavioral training and testing were declined into seven various sequences, in which Jackpot and Black Swan are either unavailable, solely or both available at a given time point of the sequence of events (Supplemental Table S1): when available, extreme events could be obtained at the 10th or the 60th activation within a given sequence, but could not occur at the same time. For example, in a behavioral sequence where the Jackpot should be available at the 10th position, any poke in the Anti-fragile and Vulnerable holes following nine responses made in one of these menus (regardless of the positive or negative outcomes) would trigger the delivery of the Jackpot. Of note, depending on the sequence used, animals could experience both extreme events in a single session.

#### Behavioral Training and Testing

Training was divided into five distinct phases before the final test: acquisition of the food collecting responses, acquisition of nose poking in the holes, training with four holes, attribution of menus to hole and training on the menus (Figure 7). Each session started with the illumination of the house light.

1. Acquisition of the food collecting responses: Animals were trained to collect sucrose pellets in the food magazine during three 30-min daily sessions (100 pellets max) under fixed ratio 1 schedule of reinforcement (FR1): one nose-poke into the food magazine triggered the delivery of one sucrose pellet. During this initial phase, nose-poke in any hole had no programmed consequences.

2. Acquisition of nose-poking in the holes: Here, animals had only access to one hole (the three others being occluded) during the 20-min sessions. One nose-poke in the available hole triggered the illumination of the hole-light and the delivery of one sucrose pellet in the food magazine. Perseverative nose-pokes (those performed before food collection) had no consequences. Following food collection in the food tray, animals were allowed to poke again in the opened hole. During each session, a maximum of 100 pellets were delivered. All animals were trained twice on each hole.

3. Training with four holes: Following the eight training sessions, animals were allowed to poke in the four different holes during twenty daily sessions of 20-min. Nose poke in any hole triggered the illumination of the associated light and the delivery of one sugar pellet in the food magazine. Both perseverative activations and pokes in other holes had no consequences. After food collection in the magazine, animals were allowed to nose poke again in any hole. The first ten training sessions were limited to 100 sugar pellets. During the following ten sessions, animals were able to collect up to 200 pellets per session.

4. Attribution of menus to hole: We determined the spatial preference for each rat by establishing the percentage of activation of each hole during the last ten sessions. To favor the emergence of Anti-fragile choices, menus’ attribution was made as follow:

• Anti-fragile exposure was associated to the preferred hole

• Robust exposure was associated to the 2nd preferred hole

• Vulnerable exposure was associated to the 3rd preferred hole

• Fragile exposure was associated to the least preferred hole

5. Training on the exposures: Here, animals were first trained on the gain domain, i.e. no time-out punishment, for each menu/hole. They were subjected to two 20-min sessions, with unlimited number of pellets, during which only one hole was available (eight training sessions in total). For Anti-fragile and Vulnerable options, we used the two sequence-types where the jackpot was available at the 10th and 60th activations (Supplemental Table S1) to ensure that all individuals could experience both an early and delayed jackpot during training. Animals were then allowed to explore all gain options (four opened holes) during 9 20-min sessions, for which different behavioral sequence-types were used. Before training on the loss domain of the different menus, animals were first exposed to a mild and constant 3-sec time-out punishment. Here, animals had access to all options, but half of the activations lead to a 3-sec time-out punishment, notified by a 3-sec tone and the extinction of the house light. During this period, pokes in the different holes or the food magazine had no programmed consequences. After the 3-sec time-out punishment, the house light was turned on and the animals could again pokes in the different holes. Following nine 20-min training sessions, the loss domain of each menus was progressively introduced. As described above, each time-out punishment was notified by a 3-sec tone and the extinction of the house light for the whole duration of the punishment, which termination was signaled by house light illumination. Animals were first exposed to concave exposures, having only access to vulnerable and fragile holes during four 20-min sessions. They were then exposed to convex losses (only Anti-fragile and Robust holes available) for another four 20-min sessions. Thus, at the end of the training, all animals experienced both extreme events at least four times.

6. Final tests: For the final tests, animals were free to explore all menus throughout the forty 20-min sessions. Animals experienced four times the ten different sequences, randomly distributed across sessions. Population (n = 20) was subdivided into two groups that experienced different, but equivalent, sequence-type distribution (Supplemental Table S2).

**Figure 7:**
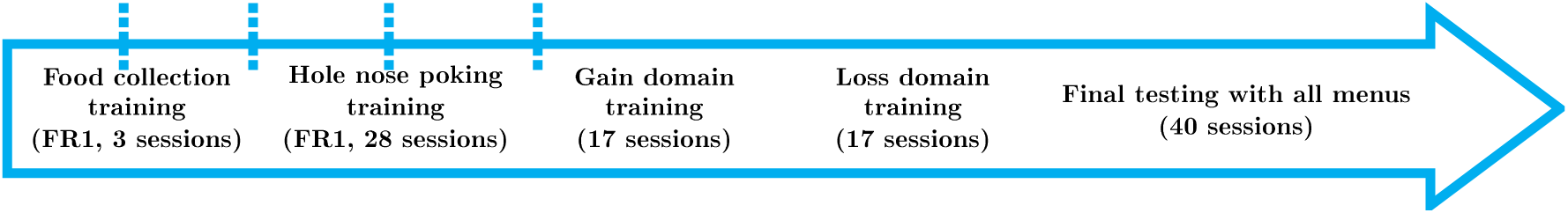
**Time-line of experiments**

Figure 7 depicts a simplified time-line of the experimental procedure.

Table S3 shows the chain of events in the ten sequences used for behavioral training and testing. Numbers in the second column indicate which event would occur and the sign preceding it whether it belongs to the gain (no sign) or loss (minus sign) domains. For example, the third event in the first sequence (noted -2) triggers a time-out punishment of 9 or 12 sec, depending on whether an animal performed his third nose poke in concave or convex menu, respectively. Note that extreme events (which should be noted 4 or -4) do not appear in the sequences, as they would automatically replace the 10th or 60th events of the sequence. See supplementary material for detailed information on sequences of events.

### Modelling and Statistical Analysis

#### Modelling Convex and Concave Exposures

Central to our experimental design is the notion of convex/concave exposure under radical uncertainty, that is, when probabilities and consequences are unknown *a priori* to subjects. A well known measure of convexity is the Jensen gap that is derived from Jensen’s inequality (see [30] for a graphical exposition), which we now define and relate to statistical moments. In our context, the relevant form of Jensen’s equality states, loosely speaking, that the expectation of a convex function of a random variable is larger than the value of that function when evaluated at the expectation of the random variable. Jensen’s gap is then defined as the difference between the expectation of the function minus the value of the function at the expectation (hence positive by construction). The inequality is reversed for a concave function and the (positive again) Jensen’s gap is then defined as the difference between the value of the function at the expectation minus the expectation of the function. In the gain domain, rats obtain 1, 3, 12 or 80 sugar pellets if the convex exposure is chosen, or 2, 4, 5, 5 pellets if the concave exposure chosen. In the loss domain, convex exposure imposes 6, 12, 15 or 15 seconds of time-out punishment, and 3, 9, 36, 240 seconds for the concave exposure. Because the values in the loss domain are proportional to the values in the gain domain, the former corresponding to 3 times the latter, we focus here on values for pellets. The statistical properties of convex and concave losses follow accordingly. More formally, sequences of gains are ordered by increasing values and are denoted 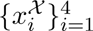 = {1, 3, 12, 80} for the convex exposure and 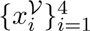 = {2, 4, 5, 5} for concave gains. We assume identical probabilities 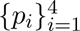 = {*p*_1_*, p*_2_*, p*_3_*, ε*} for both concave and convex gains, where *ε* is the ex-ante probability of the rare and extreme event (REE for short). The third value (that is, 12 for convex gains and 5 for concave gains) is labeled a rare event (RE), and the sets of the lowest two values (that is, 1 and 3 for convex gains, 2 and 4 for concave gains) for both exposures are composed of normal events (NE).

Making use of the exponential transform, we define Jensen gaps for the convex and concave exposures as, respectively, 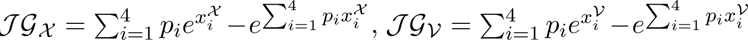, which relate to statistical moments as follows. The moment generating function for the convex and concave exposures are given by, respectively: 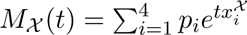 and *M_V_*(*t*) = 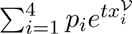. The series expansion of the exponential function allows us to derive:

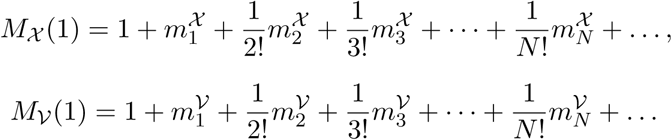

where *m^X^_N_* and *m^V^_N_* are the *N* -th statistical raw moments of the convex and concave exposures, respectively. For example, 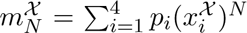 for any integer *N* ≥ 1. Jensen’s gaps for both exposures are then given by 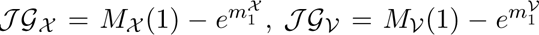. Using again the series expansion of the exponential of the first moment for each exposure, it follows that both positive Jensen gaps can be written in terms of moments, as follows:

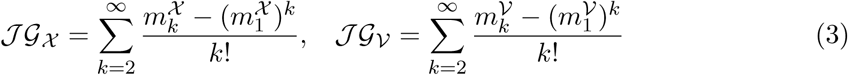

Note that the sums in equation (3) nicely allow for a straightforward decomposition in terms on all raw moments with order larger than two.

More specifically, we set probabilities to 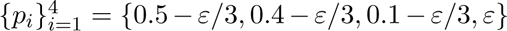 identically for both concave and convex gains. Treating the probability for the REE as a varying parameter, it can be shown that Jensen’s gaps for convex and concave exposures are ordered, with the former larger than the latter when *ε* ≤ 0.02. Over that range, it also holds true that both Jensen’s gaps are monotone increasing function of *ε*. While this property also holds for the expectations and variances of both exposures, it doesn’t for higher-order moments. For example, the skewness and kurtosis of the convex exposure turn out to be hump-shaped functions of *ε*, with peaks corresponding to values of *ε* smaller than one percent. This fact underlines that convexity measured by Jensen’s gap offers a unifying approach to rank exposures, seen as lotteries, whereas statistical moments do not necessarily do so. We conjecture that, more generally, all lotteries satisfying our assumptions (including monotone probability distribution) can be ranked according to their convexity as measured by the Jensen gap.

In Table 2, we report the central moments and the log of Jensen gaps in the top five rows, for both exposures when *ε* = 1*/*75 ≈ 1.3%, which corresponds to obtaining one REE out of 75 nose pokes. For that particular parametrization, convex and concave exposures differ mostly in their respective kurtosis and in their Jensen gaps. From row six and below, we report the decomposition of Jensen gaps in terms of the raw moments. For example, in row six is given the ratio of (*m*_2_ − (*m*_1_)^2^)*/*2 - that is, half the variance - to the corresponding Jensen gap for each exposure, in percentage. Strikingly, while moments of order 2 to 8 explain about 90% of the concave exposure’s Jensen gap, they contribute negligibly to that of the convex exposure. In fact, it turns out that moments with order around 80 start contributing, although for a small share each, to the convex exposure’s Jensen gap. In sum, while for the concave exposure a few low-order moments concentrate the contributions to the Jensen gap, the contributions of raw moments to the convex exposure’s Jensen gap spread across a larger number of very high order raw moments.

We derive next the ex-ante properties of the probability distributions associated with concave and convex gains (for definitions of stochastic dominance and congruent utility classes, see for example Fishburn and Vickson [16]; also note that first-order, resp. second-order, stochastic dominance implies second-order, resp. third-order, stochastic dominance):

- In the domain restricted to NE, concave gains first-order stochastically dominate convex gains. In addition, concave gains then have a larger expected value than that of convex gains, with equal variance, skewness and kurtosis for concave and convex exposures.
- In the domain restricted to NE and RE, concave gains second-order stochastically dominate convex gains. In addition, concave gains then have a larger expected value and smaller variance, skewness and kurtosis than that of convex gains.
- In the full domain including NE, RE and REE, convex gains second-order stochastically dominate concave gains if and only if *ε* ≥ 0.302%. In addition, convex gains then have a larger expected value, variance, skewness and kurtosis than that of concave gains if *ε* ≥ 0.302%.

The above assumptions on the experimental design are stated in terms of stochastic dominance and moments of the probability distributions. They also have implications in relation to standard approaches to decision-making. First, from an ex-ante perspective with perfect information about the probability distributions of all exposures, value-maximizing subjects whose preferences are represented by any non-decreasing value function would choose concave gains in the domain restricted to NE. They would continue to do so in the domain restricted to NE and RE for any non-decreasing and concave value function. However, in the full domain with REE, value-maximizing subjects endowed with any non-decreasing and concave value function are predicted to choose convex gains.

We denote 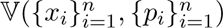 the value attributed to sequences of gains 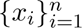 with probabilities 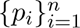, for *n* ≤ 4. The above assumptions have the following implications in terms of value maximization:

- In the domain restricted to NE, 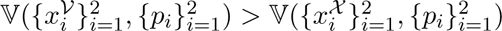 holds.
- In the domain restricted to NE and RE, 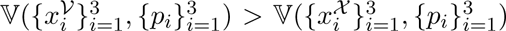 holds.
- In the unrestricted domain, 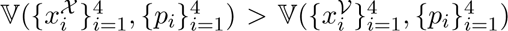 holds if and only if *ε* ≥ 0.302%.

Note that the reversal in second-order stochastic dominance (and value) in the full domain favors convex gains when extreme events that are indeed very rare - with probabilities much smaller that one percent - are added. In contrast, absent REE, adding only a RE is not enough to favor convex gains, even though it allows subjects to possibly detect convexity/acceleration and concavity/deceleration of gains and losses.

The above assumptions hold true under expected utility, that is when 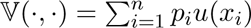 with the appropriate conditions on the utility function *u* (non-decreasing for first-order stochastic dominance, non-decreasing and concave for second-order stochastic dominance; see [16]). To better match experimental data, however, expected utility is increasingly supplemented by some form of probability weighting *w*(*p_i_*). For instance, [10] use a two-parameter functional form due to [39] and find that 30 rats out of 36 behave as if their probability weighting *w*(*p_i_*) is concave. In that case, the above ex-ante properties hold *mutatis mutandis*, for example, under expected utility with probability weighting, defined as 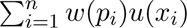. Alternatively, in the setting of rank-dependent expected utility, we can make use of results 3 and 4 in [25] to show that the above ex-ante properties still hold provided that the transformation of the cumulative distribution function is any increasing-concave function. Note that since in our experimental design REEs are both extremes (in the sense of being the largest values) and rare (that is, they have very low probabilities), one expects similar results under probability weighting and rank-dependent expected utility (or cumulative prospect theory for that matter). Although in theory assuming simply a weighting function *w*(*p_i_*) that is inverse-*s*-shaped and “very” convex for large gains (see [39] for the associated parametric restrictions) could overturn the rankings of concave and convex exposures stated above, results in [10] suggest that this is not to be expected for the overwhelming majority of the rats that are subject to their experiments.

#### Choice Data Analysis

Given the four options modelled in the previous section, each of the 20 rats went through 41 final sessions. Denote *f_A_*, *f_F_*, *f_R_*, and *f_V_* the relative frequencies (in terms of nose pokes) with which each of the four options Antifragile, Fragile, Robust and Vulnerable, respectively, is chosen over a particular session. In order to represent the rats’ choices, we construct the rotated square in panel (d) of Figure 1 using a linear transformation of the 3-frequency vector, as follows:

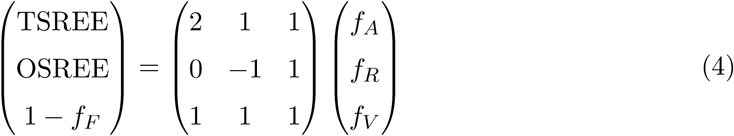

In equation (4), the first two rows deliver the two coordinates of interest that are the Total Sentivity to REE (TSREE in short) and One-sided Sentivity to REE (OSREE), while the last row requires that all frequencies sum to one. To derive TSREE and OSREE, it is useful to go through the following steps. Since we are interested in the convexity property of each chosen option, we first measure the frequencies of convex options in the gain domain by summing *f_A_* and *f_V_* . Similarly, *f_A_* + *f_R_* measures how frequent choices are convex in the loss domain. It follows that TSREE measures total convexity if it equals 2*f_A_* + *f_V_* + *f_R_*, while OSREE equals *f_V_* − *f_R_* and measures asymmetric convexity (that in the gain domain minus that in the loss domain). Hence the first two rows in equation (4). Using that *f_A_* + *f_F_* + *f_R_* + *f_V_* = 1 - that is, the last row of equation (4) - it follows that TSREE equals the simpler expression 1 + *f_A_* − *f_F_* . Note that the 3 × 3 matrix in equation (4) is invertible, which implies that all three frequencies can be recovered from the left vector. In addition, that matrix is not unique in the sense that the definitions of TSREE and OSREE that appear above as its first two lines could be moved to any other rows, provided that the identity in the third row and the constraint in the last row are adapted accordingly. This essentially means that given both differences TSREE (= 1 + *f_A_* − *f_F_*) and OSREE (= *f_V_* − *f_R_*), any of the four relative frequencies provides enough information to derive the remaining three, using the constraint that all four frequencies sum up to one. Finally, it follows that in the rotated square in panel (d) of Figure 1, the four vertices have the following coordinates: (0, 0) for Fragile, (0, 2) for Antifragile, (−1, 1) for Robust, and (1, 1) for Vulnerable. TSREE and OSREE are depicted in Figure 2.

TSREE and OSREE are also related to behavioral measures of the share of nosepokes that lead to either exposure or non exposure to REE, as follows: we define Black Swan Avoidance as the share of nosepokes that lead to avoidance of Black Swans - i.e. *f_A_* + *f_R_* - and Jackpot Seeking as the share of nosepokes that lead to be exposed to Jackpots - i.e. *f_A_* + *f_V_* . It follows that TSREE is by definition the sum of those two shares, while OSREE is the difference between the former and the latter. Formally, if we denote JPS and BSA the shares of nosepokes that lead to be exposed to Jackpots and to avoid Black Swans, this implies that JPS= 0.5(TSREE+OSREE) and BSA= 0.5(TSREE−OSREE). In other words, in the (not rotated) square that appears in panel (d) of Figure 1, the four vertices have the alternative coordinates: (0, 0) for Fragile, (1, 1) for Antifragile, (0, 1) for Robust, and (1, 0) for Vulnerable, in the (JPS,BSA) coordinates. Obviously, both interpretations are equivalent up to a change in coordinates which is a bijection. Note that in the rotated square in panel (d) of Figure 1, lines with a 45-degree slope depict choices with constant BSA so that lines moving north-west represent increasing BSA. Similarly, lines with a −45-degree slope depict choices with constant JPS so that lines moving north-east represent increasing JPS. Median BSA and JP are depicted in Figure 3.

In Figure 4 we report the short term behavioral responsiveness of rat to REE. Behavioral responsiveness is measured by calculating for each rat before/after differences in TSREE and OSREE. This amounts to calculate TSREE and OSREE over the 10 choices preceding a REE and over the 10 choices following the occurrence of each REE. The window of 10 choices before and after is dictated by the configuration of REE in our design. A REE can happen at 11th choice in a sequence (see appendix). Before/after TSREE and OSREE are averaged by rat over the 41 sessions in the gain domain (Jackpot) and in the loss domain (Black Swan). Table 3 reports the values for the mean before/after differences. In each panel of Figure 4, each rat is depicted by a color-coded dot and the dotted black line represents the 45-degree line. Color-coded dot on the 45-degree line indicate no difference in TSREE or OSREE before and after a REE, i.e. no short term behavioral responsiveness to REE. To visualisation purpose, we computed for each panel a smooth spline regression that estimates a non parametric relationship between before and after TSREE and OSREE. They are plotted as solid black lines in each panel.

We then test the statistical significance of short term behavioral by conducting bootstrap paired-sample mean tests on before/after coordinates (separately) for each type of REE. We assessed short-term behavioral responses using a paired bootstrap test (boot.paired.bca) in R (wBoot package). This non-parametric method estimates the mean difference between paired conditions by resampling with replacement and computes a bias-corrected and accelerated (BCa) confidence interval, providing robust inference without assuming normality. An observation in the sample is the mean behavioral responsiveness in TSREE or OSREE for a given rat. Observations therefore respect statistical independence. Bootstrap tests lead to the following *p*-values, under the null hypothesis that the mean difference is zero. Following Jackpots, *p* = 0.3470 for TSREE and *p* = 0.0522 for OSREE, which indicates that the null hypothesis is not rejected at 1%. Following Black Swans, however, the null hypothesis is rejected at 1% since *p* = 0.0045 for TSREE and *p* = 0.0064 for OSREE. In sum, the mean before/after difference following a REE in the loss domain is significant for both Total and One-sided sensitivities, further confirming the Black Swan avoidance result that we document in this paper.

We also report in Figure 8 how choices by rats after the 41 final sessions compare to choices made in the training step 3, defined in the section on Behavioral Training and Testing. We do so by computing TSREE and OSREE in step 3 sessions and pair coordinates resulting from training with that resulting from the 40 experimental sessions reported in Figure 2. The lines between two color-coded dot connects for each rat its conditioning coordinates with its coordinates resulting from its behavior in the 41 sessions. We report on the right side of the figure the total variation distance between the distribution of nosepokes in the 4 holes in the training and the distribution in the 41 experimental sessions. The interpretation of total variation distance in our setting is simple: it measures, for each rat, the proportion of nosepokes that need to be changed in order to equalize the behavior in the training and the behavior in the 41 final experimental sessions. Median and mean total variation distance are 0.293 and 0.324 respectively, which implies that half of the rats changed more than 30% of their choices in the final experimental sessions compared to the training sessions.

**Figure 8:**
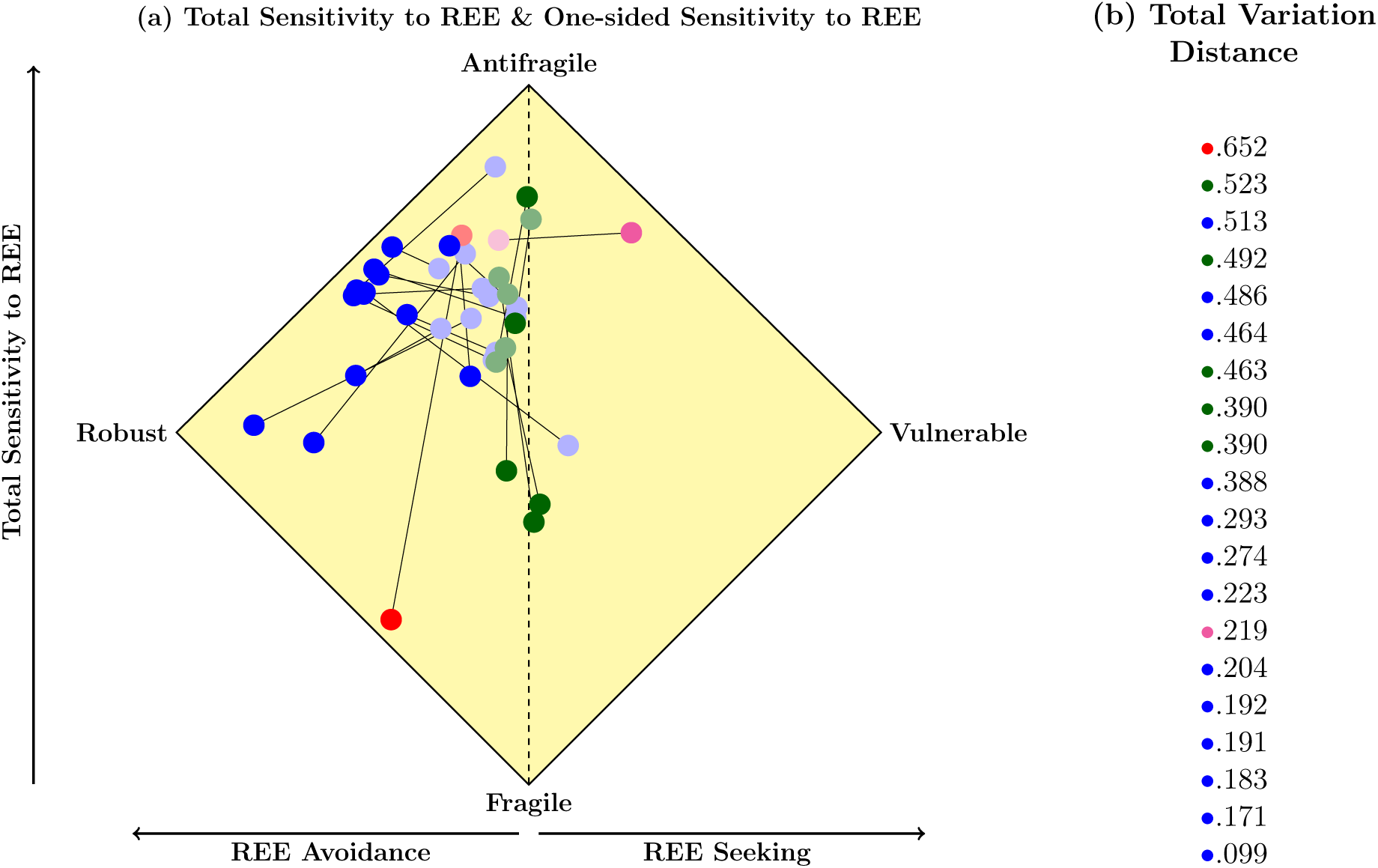
Variation of choices observed in final experimental sessions compared to that observed in training sessions (step 3 in Material and Methods section on Experimental Method Details); panel (a) shows variations of Total Sensitivity to REE (*y*-axis) and of One-sided Sensitivity to REE (*x*-axis), that is, dark color-coded dots replicate averages over the 40 sessions for each of the 20 rats - see panel (c) in Figure 2 - and are paired with averages over training sessions for the same rat, depicted with light color-coded dots connected through lines with corresponding dark color-coded dots; panel (b) shows total variation distance, that is, the proportion of total nose pokes in training sessions that need to be changed to replicate nosepokes in final sessions

Finally, in Figure S1 of the Supplementary Material, we report what we label “convexity premiums”, which are counterfactual situations that we construct as follows. Given the rats’ choices over the 41 sessions, we recall all random sequences that have been used to generate gains and losses for each rat. We next compute the outcome that would have been obtained, had the rat chosen convex options in the gain domain (that is, Antifragile or Vulnerable) and in the loss domain (that is, Antifragile or Robust) all the time. The right and left panels in Figure S1 depict the normalized convexity premiums thus computed, for each rat represented in rows, with convex exposures yielding more pellets and implying less waiting in terms of seconds.

#### Augmented Q-Learning Model Estimation and Simulation

Parameter optimization is carried out for each rat over the 41 sessions, using observed choices and outcomes. We estimate by maximum likelihood two series of augmented Q-Learning models. The first series integrates REE in Q subvalues whereas the second series does not. Each series consists of a first baseline model without specific forgetting rates and decision weights attached to REE. Two partial models integrate either specific forgetting parameters or decisions weights on the presence of REE. The fourth model integrate both. In total, we estimate eight nested models for each rat. Formally, we estimate four different models: a baseline model (Model 1) without specific forgetting parameters (*α^f^_k_* = *α_k_*, with *k* ∈ {*g, l*}) and with zero decision weights of REE (*γ_g_*= *γ_l_* = 0). Model 2 introduces specific forgetting parameters (*α^f^_k_* = *α_k_*, with *k* ∈ {*g, l*}) and Model 3 introduces decision weights of REE (*γ_g_* and *γ_l_* ≠ 0). Model 4 introduces both.

To ensure convergence and avoid local maxima, each parameter in baseline models was initialized with two different points of its parameter space. This makes in total 4^2^ = 16 combinations of initial parameter values. Each of the 16 models is estimated using an automatized two-step procedure. The fist step consists of an unconstrained downhill simplex method in [31], which does not make use of first derivatives. Estimated parameters are then used as initial parameters in the second step of the estimation procedure, a quasi-Newton method in [17]. Parameter estimates from baseline models are then used as initial parameters for subsequent augmented models. Estimation is carried out using the above two-step procedure to ensure that we convergence is reached.

Once the eight models are estimated, we then compare them using both Akaike information criterion (AIC) and Bayesien Information criterion (BIC), that penalizes the use of additional parameters, so as to select which model configuration best fits observed behavior for each rat (see Supplementary Material for detailed results per rat). We check whether the selected model properly estimates sensitivities to REE by comparing estimated sensitivities to observed sensitivities to REE. This is done by computing Total Sensitivity to REE and One-sided Sensitivity to REE based on estimated choice frequencies from the extended Q-Learning model over the 41 sessions. Fitted sensitivities are reported bottom right panel in Figure 5. Finally, we checked that selected models were able to reproduce experimental data by running simulations for each phenotype observed in the sample of animals (as in [35]). Simulations are conducted over the artificially generated original 41 sessions, using individual parameter estimates for pink and red rats, median parameter estimates for blue and green rats and estimated Q subvalues as prior Qs. For each simulation, we then computed Total Sensitivity to REE and One-sided Sensitivity to REE. Simulations are reported in bottom right panel of Figure 5. See Supplementary Material for more details on the estimation results and for an exploration in related outcome range-adaption models.

## Acknowledgments

Funding: The authors are grateful for the support of ANR through BEAM (ANR-15-ORAR-0004-03) and of CNRS through a MITI interdisciplinary grant (call for projects on rare events).

## Author contributions

C.B., M.D, S.L., and P.P. designed the experiments. M.D and L.W.M. performed the experiments. S.L. and P.P. analyzed and modelled the data. S.L, P.P. and C.B, wrote the manuscript.

## Competing interests

The authors declare that they have no competing interests.

## Data and materials availability

All data needed to replicate the results are available via GitHub at https://github.com/PatrickPintus/Rare-and-Extreme-Events. Code is available from the authors upon request .

## Supplementary Material

### Sequences used in the experimental design

In Table S1 we report the different types of sequences used for behavioral training and testing, which shows the position of the extreme events. For example, in the sequence-type 6, jackpot could be obtained at the 10th position while the Black-swan would be triggered at the 60th activation.

**Table S1:** Position of REE in the sequence.

| # | Jackpot | Black Swan |
| --- | --- | --- |
| 1 |  |  |
| 2 | 10 |  |
| 3 | 60 |  |
| 4 |  | 10 |
| 5 |  | 60 |
| 6 | 10 | 60 |
| 7 | 60 | 10 |

Table S2 shows the different behavioral sequence-types used for the forty testing sessions. As indicated in the text, half of the population was subjected to the sequence-types described in the left part of the ‘type column’, while the other half experienced the sequence-types described in the right part of the ‘type column’. Table S3 shows the succession of events for each sequence.

**Table S2:** Sequences and types.

| Session | Sequence | Type |  |
| --- | --- | --- | --- |
| 1 | 2 | 5 | 3 |
| 2 | 10 | 6 | 5 |
| 3 | 5 | 1 | 3 |
| 4 | 3 | 7 | 1 |
| 5 | 4 | 3 | 3 |
| 6 | 1 | 7 | 6 |
| 7 | 6 | 2 | 6 |
| 8 | 6 | 6 | 2 |
| 9 | 4 | 2 | 4 |
| 10 | 1 | 1 | 1 |
| 11 | 7 | 7 | 4 |
| 12 | 1 | 3 | 2 |
| 13 | 7 | 7 | 7 |
| 14 | 8 | 2 | 4 |
| 15 | 5 | 2 | 5 |
| 16 | 7 | 2 | 6 |
| 17 | 3 | 1 | 6 |
| 18 | 5 | 5 | 4 |
| 19 | 2 | 1 | 6 |
| 20 | 7 | 6 | 1 |
| 21 | 9 | 4 | 4 |
| 22 | 4 | 2 | 6 |
| 23 | 2 | 5 | 2 |
| 24 | 2 | 3 | 5 |
| 25 | 4 | 2 | 2 |
| 26 | 9 | 2 | 2 |
| 27 | 10 | 5 | 6 |
| 28 | 10 | 5 | 4 |
| 29 | 10 | 1 | 4 |
| 30 | 3 | 3 | 6 |
| 31 | 8 | 5 | 4 |
| 32 | 3 | 7 | 6 |
| 33 | 6 | 2 | 1 |
| 34 | 9 | 4 | 4 |
| 35 | 9 | 3 | 5 |
| 36 | 6 | 5 | 1 |
| 37 | 1 | 5 | 3 |
| 38 | 8 | 4 | 4 |
| 39 | 5 | 3 | 7 |
| 40 | 8 | 4 | 4 |

**Table S3:**
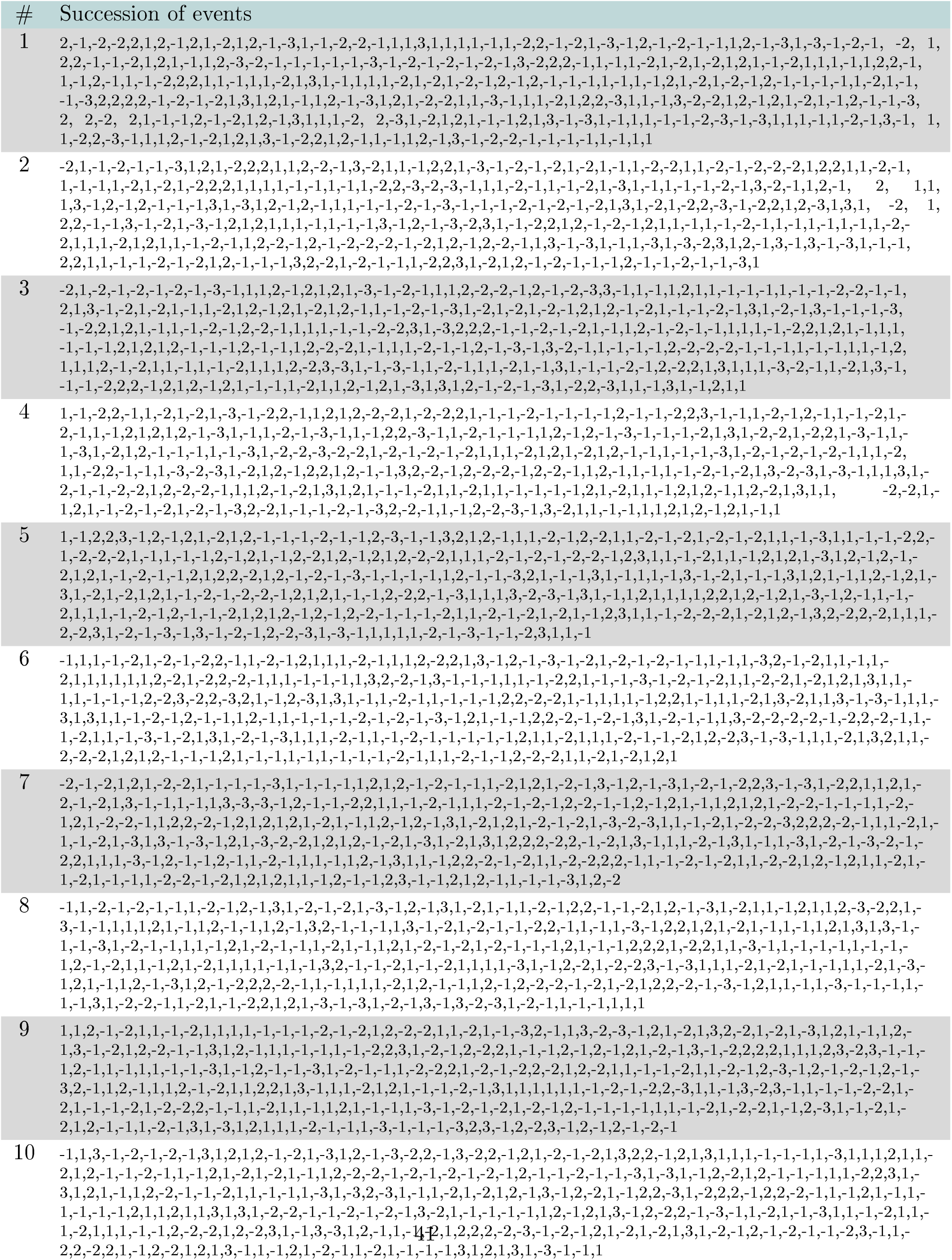
Succession of events for each sequence.

### Behavioral measures for each rat

All behavioral measures for each rat that are used in the main text are gathered in Table S4: Total and One-sided Sensitivities to REE, Black Swan Avoidance and Jackpot Seeking (both in %).

Table S4: Behavioral measures for each rat - Total and One-sided Sensitivities to REE - TSREE and OSREE in short; Black Swan Avoidance and Jackpot Seeking in percentage - BSA and JPS in short. The last four columns present the percentage choice of each option.

**Table S4:** Behavioral measures for each rat - Total and One-sided Sensitivities to REE - TSREE and OSREE in short; Black Swan Avoidance and Jackpot Seeking in percentage - BSA and JPS in short. The last four columns present the percentage choice of each option.

| Rat | TSREE | OSREE | BSA | JPS | Anti-fragile | Robust | Vulnerable | Fragile |
| --- | --- | --- | --- | --- | --- | --- | --- | --- |
| • 1 | 1.4136 | -0.4583 | 93.59 | 47.76 | 44.76 | 49.69 | 03.93 | 01.61 |
| • 2 | 1.2177 | -0.1392 | 67.84 | 53.92 | 44.27 | 22.82 | 06.68 | 26.21 |
| • 3 | 0.9749 | -0.5808 | 77.78 | 19.70 | 09.08 | 70.11 | 09.60 | 11.20 |
| • 4 | 1.2264 | -0.4259 | 82.61 | 40.02 | 43.66 | 52.21 | 01.68 | 02.43 |
| • 5 | 1.4208 | -0.4990 | 95.99 | 46.09 | 45.36 | 49.11 | 01.43 | 04.07 |
| • 6 | 1.0819 | -0.6024 | 84.21 | 23.97 | 10.57 | 80.32 | 01.80 | 07.28 |
| • 7 | 0.4632 | 0.0122 | 22.55 | 23.77 | 35.50 | 02.49 | 03.98 | 58.02 |
| • 8 | 1.3706 | -0.0669 | 71.87 | 65.18 | 17.39 | 13.46 | 17.34 | 51.79 |
| • 9 | 0.6928 | 0.0304 | 33.12 | 36.16 | 32.89 | 05.67 | 09.02 | 52.40 |
| • 10 | 1.4652 | -0.4058 | 93.55 | 52.97 | 49.49 | 45.49 | 02.73 | 02.27 |
| • 11 | 0.9074 | 0.0104 | 44.85 | 45.89 | 34.67 | 12.53 | 05.80 | 46.99 |
| • 12 | 1.4365 | -0.2772 | 85.68 | 57.96 | 47.79 | 37.46 | 02.26 | 12.47 |
| • 13 | 1.5444 | -0.1653 | 85.48 | 68.95 | 58.94 | 31.91 | 09.13 | 00.00 |
| • 14 | 1.5504 | -0.3937 | 97.20 | 57.83 | 56.61 | 40.59 | 01.28 | 01.50 |
| • 15 | 1.6610 | 0.0447 | 80.81 | 85.28 | 75.86 | 08.43 | 08.52 | 07.17 |
| • 16 | 1.4076 | -0.5050 | 95.63 | 45.13 | 43.56 | 52.04 | 01.59 | 02.79 |
| • 17 | 0.5661 | -0.3929 | 47.95 | 08.66 | 03.10 | 40.79 | 01.10 | 54.99 |
| • 18 | 1.1406 | -0.4850 | 81.28 | 32.78 | 30.14 | 53.34 | 03.68 | 12.82 |
| • 19 | 1.6009 | 0.2997 | 65.06 | 95.03 | 61.61 | 02.96 | 32.94 | 02.46 |
| • 20 | 1.4832 | -0.4491 | 96.61 | 51.70 | 51.09 | 45.52 | 00.88 | 02.49 |

### Augmented Q-Learning model selection according to BIC

Table S5 presents BIC values for all rat models. In the first column, the rats’ phenotypes are color-coded as in the text. Each uniquely selected model is identified by light blue color.

**Table S5:** BIC values for all augmented Q-Learning models, with selected value highlighted.

|  | REE in Q-subvalues |  |  |  | REE not in Q-subvalues |  |  |  |
| --- | --- | --- | --- | --- | --- | --- | --- | --- |
|  | Model 1 | Model 2 | Model 3 | Model 4 | Model 1 | Model 2 | Model 3 | Model 4 |
| • 1 | 10147.618 | 9473.977 | 9455.796 | 9233.393 | 9639.117 | 8909.843 | 9048.188 | 8883.051 |
| • 2 | 13215.452 | 13188.041 | 13121.341 | 13135.077 | 13061.968 | 13040.253 | 13037.390 | 13042.841 |
| • 3 | 8825.413 | 8748.519 | 8832.939 | 8754.428 | 8668.958 | 8581.156 | 8679.882 | 8592.356 |
| • 4 | 9559.508 | 9569.403 | 9118.611 | 9127.721 | 9818.665 | 9473.840 | 9338.137 | 9355.298 |
| • 5 | 8851.442 | 8837.212 | 8737.694 | 8753.888 | 8842.836 | 8831.712 | 8728.778 | 8743.304 |
| • 6 | 7206.668 | 7026.401 | 7058.036 | 7040.684 | 7113.624 | 6824.310 | 6931.265 | 6812.231 |
| • 7 | 10194.305 | 9407.034 | 9817.474 | 9419.479 | 9659.679 | 9184.589 | 9559.764 | 9135.728 |
| • 8 | 12495.756 | 12506.051 | 12505.885 | 12515.155 | 12492.775 | 12496.156 | 12490.187 | 12489.188 |
| • 9 | 12613.911 | 12183.914 | 12581.295 | 12187.231 | 12194.450 | 11855.559 | 12195.525 | 11831.083 |
| • 10 | 8822.488 | 8532.439 | 8612.817 | 8624.451 | 8269.455 | 8130.544 | 8083.416 | 8018.394 |
| • 11 | 11119.104 | 11005.450 | 11006.259 | 10950.474 | 11030.692 | 10873.231 | 10981.489 | 10889.932 |
| • 12 | 11387.450 | 11198.078 | 11105.750 | 11069.321 | 11222.624 | 11032.712 | 11089.737 | 11065.547 |
| • 13 | 10835.134 | 10296.487 | 10388.582 | 10241.068 | 10677.363 | 10215.775 | 10373.480 | 10202.434 |
| • 14 | 10210.664 | 9855.573 | 9536.958 | 9552.319 | 9996.584 | 9880.569 | 9521.883 | 9305.535 |
| • 15 | 9042.715 | 8933.762 | 9051.019 | 8895.580 | 8964.300 | 8840.706 | 8981.304 | 8824.601 |
| • 16 | 10395.829 | 10219.158 | 9990.742 | 10003.734 | 10600.320 | 10107.624 | 10019.103 | 10035.935 |
| • 17 | 9636.231 | 9191.330 | 9164.951 | 9152.413 | 9661.941 | 9285.094 | 9126.409 | 9143.220 |
| • 18 | 12948.075 | 12737.650 | 12777.029 | 12685.432 | 12808.119 | 12537.832 | 12676.942 | 12523.725 |
| • 19 | 11146.154 | 10803.117 | 10362.629 | 10371.896 | 11347.877 | 10585.857 | 10230.564 | 10231.779 |
| • 20 | 9885.053 | 9359.053 | 9279.145 | 9293.579 | 9742.212 | 9304.446 | 9292.413 | 9307.507 |

### Augmented Q-Learning model selection according to AIC

Table S6 presents AIC values for all rat models. In the first column, the rats’ phenotypes are color-coded as in the text. Each uniquely selected model is identified by light blue color.

**Table S6:** AIC values for all augmented Q-Learning models, with selected value highlighted.

|  | REE in Q-subvalues |  |  |  | REE not in Q-subvalues |  |  |  |
| --- | --- | --- | --- | --- | --- | --- | --- | --- |
|  | Model 1 | Model 2 | Model 3 | Model 4 | Model 1 | Model 2 | Model 3 | Model 4 |
| • 1 | 10120.983 | 9434.024 | 9415.843 | 9180.122 | 9612.482 | 8869.889 | 9008.235 | 8829.780 |
| • 2 | 13188.983 | 13148.337 | 13081.638 | 13082.139 | 13035.499 | 13000.550 | 12997.686 | 12989.903 |
| • 3 | 8799.339 | 8709.409 | 8793.829 | 8702.281 | 8642.884 | 8542.045 | 8640.772 | 8540.209 |
| • 4 | 9533.087 | 9529.770 | 9078.979 | 9074.878 | 9792.244 | 9434.208 | 9298.504 | 9302.455 |
| • 5 | 8825.399 | 8798.149 | 8698.630 | 8701.804 | 8816.794 | 8792.648 | 8689.714 | 8691.220 |
| • 6 | 7180.386 | 6986.979 | 7018.614 | 6988.121 | 7087.343 | 6784.888 | 6891.842 | 6759.669 |
| • 7 | 10167.856 | 9367.360 | 9777.800 | 9366.581 | 9633.230 | 9144.915 | 9520.090 | 9082.829 |
| • 8 | 12469.528 | 12466.708 | 12466.542 | 12462.697 | 12466.546 | 12456.813 | 12450.844 | 12436.730 |
| • 9 | 12587.291 | 12143.983 | 12541.364 | 12133.989 | 12167.829 | 11815.628 | 12155.594 | 11777.841 |
| • 10 | 8796.092 | 8492.845 | 8573.222 | 8571.658 | 8243.059 | 8090.949 | 8043.821 | 7965.600 |
| • 11 | 11093.143 | 10966.508 | 10967.317 | 10898.552 | 11004.731 | 10834.290 | 10942.548 | 10838.011 |
| • 12 | 11361.251 | 11158.780 | 11066.452 | 11016.924 | 11196.425 | 10993.414 | 11050.439 | 11013.149 |
| • 13 | 10808.770 | 10256.941 | 10349.036 | 10188.340 | 10650.999 | 10176.229 | 10333.934 | 10149.706 |
| • 14 | 10183.890 | 9815.412 | 9496.797 | 9498.771 | 9969.810 | 9840.408 | 9481.722 | 9251.987 |
| • 15 | 9016.017 | 8893.716 | 9010.973 | 8842.185 | 8937.603 | 8800.660 | 8941.258 | 8771.206 |
| • 16 | 10369.190 | 10179.198 | 9950.783 | 9950.455 | 10573.680 | 10067.665 | 9979.144 | 9982.656 |
| • 17 | 9609.738 | 9151.590 | 9125.210 | 9099.426 | 9635.448 | 9245.354 | 9086.669 | 9090.233 |
| • 18 | 12921.281 | 12697.459 | 12736.838 | 12631.843 | 12781.325 | 12497.641 | 12636.751 | 12470.136 |
| • 19 | 11119.323 | 10762.871 | 10322.383 | 10318.235 | 11321.046 | 10545.611 | 10190.318 | 10178.118 |
| • 20 | 9858.545 | 9319.291 | 9239.382 | 9240.562 | 9715.703 | 9264.684 | 9252.650 | 9254.490 |

### Parameter estimates for selected Q-Learning models

Table S7 presents for each rat all estimated parameters arising from models selected using BIC (see Table S5 for values of this criterion).

**Table S7:** Parameter estimates by rat for all selected Q-Learning models.

| Rat | $\lambda_g$ | $\lambda_l$ | $\alpha_g$ | $\alpha_l$ | $\gamma_g$ | $\gamma_l$ | $\alpha_g^f$ | $\alpha_l^f$ |
| --- | --- | --- | --- | --- | --- | --- | --- | --- |
| • 1 | 1.8905 | -0.2989 | 0.2167 | 0.0186 | 0.1326 | -0.9778 | 1.0000 | 0.0023 |
| • 2 | 2.5173 | -1.9655 | 0.0095 | 0.0001 | -0.2132 | -0.1046 | 0.0095 | 0.0001 |
| • 3 | 10.2292 | -0.4393 | 0.0242 | 0.0172 | 0.0000 | 0.0000 | 1.0000 | 0.0084 |
| • 4 | 0.2694 | -0.5987 | 0.8493 | 0.0060 | -0.1491 | -0.1276 | 0.8493 | 0.0060 |
| • 5 | 0.0853 | -1.3210 | 1.0000 | 0.0000 | -0.1456 | -3.0508 | 1.0000 | 0.0000 |
| • 6 | 2.9762 | -0.4528 | 0.0884 | 0.0210 | 0.5453 | -0.1583 | 1.0000 | 0.0028 |
| • 7 | 0.8926 | -0.5457 | 0.2932 | 0.0058 | -0.4648 | 0.2237 | 1.0000 | 0.0001 |
| • 8 | 0.5390 | -0.2567 | 0.0474 | 0.0147 | -0.0042 | -0.2049 | 0.0325 | 0.0033 |
| • 9 | 11.7451 | -0.3810 | 0.0216 | 0.0088 | -0.1397 | 0.1999 | 1.0000 | 0.0008 |
| • 10 | 1.0972 | -0.1673 | 0.6405 | 0.0412 | 0.1963 | -1.6800 | 1.0000 | 0.0001 |
| • 11 | 1.0574 | -0.1694 | 0.0124 | 0.0509 | 0.0000 | 0.0000 | 0.0016 | 0.0047 |
| • 12 | 0.8191 | -0.2108 | 0.0047 | 0.0510 | 0.0000 | 0.0000 | 0.0002 | 0.0054 |
| • 13 | 2.8084 | -0.2902 | 0.0045 | 0.0534 | 0.1628 | -0.5758 | 0.0275 | 0.0019 |
| • 14 | 0.4545 | -0.0711 | 0.3508 | 0.6407 | 0.2730 | -3.2954 | 1.0000 | 0.8990 |
| • 15 | 0.5887 | -0.3196 | 0.0316 | 0.0439 | 0.0377 | 0.0938 | 0.0017 | 0.0062 |
| • 16 | 0.0364 | -0.1964 | 0.0003 | 0.0308 | -0.1129 | -2.6553 | 0.0003 | 0.0308 |
| • 17 | 0.3238 | -4.0915 | 0.7455 | 0.0000 | -2.6254 | 0.1739 | 0.7455 | 0.0000 |
| • 18 | -0.3281 | -0.3422 | 0.5824 | 0.0301 | -0.2130 | -0.3513 | 1.0000 | 0.0033 |
| • 19 | 1.7988 | 0.0548 | 0.0001 | 0.9999 | 2.7104 | -0.5323 | 0.0001 | 0.9999 |
| • 20 | 2.0923 | 40.1887 | 0.0009 | 0.0000 | 0.0746 | -3.5294 | 0.0009 | 0.0000 |

### Comparison of mean and median parameter estimates of augmented Q-Learning models for blue and green phenotypes

Table S8 presents median and mean parameter values estimated from augmented Q-Learning models for green and blue phenotypes, as well as the relevant statistical tests. Parameters difference tests are carried out by Wilcoxon Rank Sum tests (column 4) and Mean Difference tests (column 7). Significant results are highlighted.

**Table S8:** Mean and median parameter estimates for blue and green phenotypes, and difference tests with significant results highlighted.

| Parameter | Median<br>blue<br>rats (●) | Median<br>green<br>rats (●) | Wilcoxon<br>Rank Sum<br>test (p-value) | Mean<br>blue<br>rats (●) | Mean<br>green<br>rats (●) | Mean<br>Difference<br>test (p-value) |
| --- | --- | --- | --- | --- | --- | --- |
| $\lambda_g$ | 1.0972 | 0.8926 | 0.8490 | 1.9217 | 2.9646 | 0.6717 |
| $\lambda_l$ | -0.2989 | -0.3196 | 1.0000 | 1.9834 | -0.3345 | 0.3674 |
| $\gamma_g$ | 0.0000 | -0.0042 | 0.3233 | 0.0425 | -0.1142 | 0.1973 |
| $\gamma_l$ | -0.5758 | 0.0938 | 0.0102 | -1.2700 | 0.0625 | 0.0047 |
| $\alpha_g$ | 0.0884 | 0.0316 | 0.8490 | 0.2902 | 0.0813 | 0.2126 |
| $\alpha_l$ | 0.0210 | 0.0147 | 0.9241 | 0.0700 | 0.0248 | 0.9694 |
| $\alpha_g^f$ | 1.0000 | 0.0325 | 0.6331 | 0.6067 | 0.4072 | 0.2875 |
| $\alpha_l^f$ | 0.0028 | 0.0033 | 0.9241 | 0.0739 | 0.0030 | 0.4526 |

### An augmented Q-Learning model with outcome range-adaptation

As a robustness check, we introduce in our augmented Q-learning models the range principle due to [36] - see also [34] - which captures the notion that subjective evaluation of rewards may take into account their range through the Min-Max normalization presented below. That is, we replace objective rewards by their subjective judgements *s*(.) as follows:

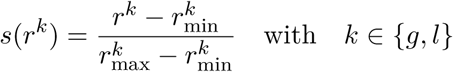

Subjective judgements were then substituted in equation (1) for gains and losses and the resulting Q-learning models were estimated using the procedure described in Material and Methods, section Augmented Q-Learning Model Estimation and Simulation. More precisely, subjective judgements for gains and losses are as follows: *s*(*r^g^*) = (*r^g^* − 1)*/*(80 − 1) for options that give convex gains and *s*(*r^g^*) = (*r^g^*− 2)*/*(5 − 2) for options that give concave gains; similarly, *s*(*r^l^*) = (*r^l^*+ 15)*/*(−6 + 15) when losses are convex losses and *s*(*r^l^*) = (*r^l^* + 240)*/*(−3 + 240) when losses are concave. REE are then used (as Min and Max values) to normalize all rewards and they are included in the Q-sub-values when they happen. This implies, by definition, that subjective judgements equal zero when Black Swans materialize, and equal one when Jackpots occur.

When REE are introduced through the normalization of gains and losses described above, models that consider nonzero decisions weights for REE are selected for 18 rats out of 20. Augmented Q-learning models with outcome range-adaptation proved however to improve the quality of the models for four rats only and we kept with our parsimonious models in the article. BIC and AIC values by rat are presented in the Tables S9-S10, for all four models with REE included in the Q-sub-values.

**Table S9:** BIC values for all augmented Q-Learning model with outcome range-adaptation, with selected value highlighted.

|  | Model 1 | Model 2 | Model 3 | Model 4 |
| --- | --- | --- | --- | --- |
| • 1 | 10360.052 | 9731.387 | 9632.944 | 9497.435 |
| • 2 | 13380.594 | 13389.451 | 13101.613 | 13021.902 |
| • 3 | 8867.516 | 8755.716 | 8863.804 | 8750.195 |
| • 4 | 9668.138 | 9630.168 | 9388.315 | 9404.728 |
| • 5 | 8974.453 | 8937.020 | 8730.574 | 8746.844 |
| • 6 | 7323.166 | 7081.788 | 7116.992 | 6967.601 |
| • 7 | 9921.860 | 9584.036 | 9932.662 | 9542.869 |
| • 8 | 12817.763 | 12788.346 | 12639.371 | 12656.187 |
| • 9 | 12750.633 | 12648.171 | 12219.681 | 11858.015 |
| • 10 | 9565.135 | 9199.293 | 9316.995 | 9067.288 |
| • 11 | 11299.508 | 11169.590 | 11093.347 | 10943.598 |
| • 12 | 11255.261 | 11141.949 | 11103.222 | 11014.454 |
| • 13 | 10563.680 | 10416.064 | 10282.824 | 10197.848 |
| • 14 | 10072.404 | 9837.106 | 9499.525 | 9497.333 |
| • 15 | 8921.307 | 8861.083 | 8865.852 | 8868.116 |
| • 16 | 10637.961 | 10207.127 | 9981.039 | 9997.678 |
| • 17 | 9803.718 | 9282.518 | 9303.068 | 9305.074 |
| • 18 | 12827.397 | 12787.223 | 12602.462 | 12505.592 |
| • 19 | 10673.745 | 10668.086 | 10371.852 | 10387.190 |
| • 20 | 9796.662 | 9334.566 | 9295.024 | 9261.452 |

**Table S10:** AIC values for all augmented Q-Learning model with outcome range-adaptation, with selected value highlighted.

|  | Model 1 | Model 2 | Model 3 | Model 4 |
| --- | --- | --- | --- | --- |
| • 1 | 10333.417 | 9691.434 | 9592.991 | 9444.164 |
| • 2 | 13354.126 | 13349.747 | 13061.910 | 12968.964 |
| • 3 | 8841.442 | 8716.605 | 8824.694 | 8698.048 |
| • 4 | 9641.717 | 9590.536 | 9348.683 | 9351.885 |
| • 5 | 8948.411 | 8897.957 | 8691.511 | 8694.759 |
| • 6 | 7296.884 | 7042.366 | 7077.570 | 6915.038 |
| • 7 | 9895.410 | 9544.362 | 9892.988 | 9489.970 |
| • 8 | 12791.534 | 12749.003 | 12600.028 | 12603.729 |
| • 9 | 12724.012 | 12608.240 | 12179.750 | 11804.773 |
| • 10 | 9538.739 | 9159.698 | 9277.400 | 9014.495 |
| • 11 | 11273.547 | 11130.648 | 11054.406 | 10891.676 |
| • 12 | 11229.062 | 11102.651 | 11063.924 | 10962.057 |
| • 13 | 10537.316 | 10376.518 | 10243.278 | 10145.120 |
| • 14 | 10045.630 | 9796.945 | 9459.364 | 9443.785 |
| • 15 | 8894.609 | 8821.036 | 8825.806 | 8814.721 |
| • 16 | 10611.321 | 10167.168 | 9941.080 | 9944.399 |
| • 17 | 9777.224 | 9242.777 | 9263.328 | 9252.088 |
| • 18 | 12800.603 | 12747.032 | 12562.271 | 12452.004 |
| • 19 | 10646.914 | 10627.840 | 10331.606 | 10333.529 |
| • 20 | 9770.154 | 9294.803 | 9255.261 | 9208.435 |

### Outcomes

What are the ex-post outcomes of all choices made by the different rats over the course of the 41 final sessions? In the first left columns of Table S11, we report for each rat the number of nose pokes, the number of REE of each type the rat has experienced. We also report, for the rewards in pellets and the waiting times in seconds, the sum as well as the first four moments of the outcome per nose poke. The overall pattern that emerges from Table S11 is that a typical rats in with high Total Sensitivity group tends to have outcomes that differ from a typical rat with moderate Total Sensitivity. For gains (i.e. sugar pellets), the former’s rewards have typically higher mean and variance but smaller skewness and kurtosis that the later’s. This happens for instance, to an extreme degree, when we compare rat 15 (the most Anti-fragile rat) and rat 17. On the loss side, by symmetry the waiting time of the average high-sensitivity rat tends to have lower mean and variance, but larger skewness and kurtosis. These facts are consistent with the fact that a typical high-Total Sensitivity rat tend to pick a mix of exposures that is more convex both on the gain domain and on the loss domain, compared to an average low-Total Sensitivity rat.

**Table S11:**
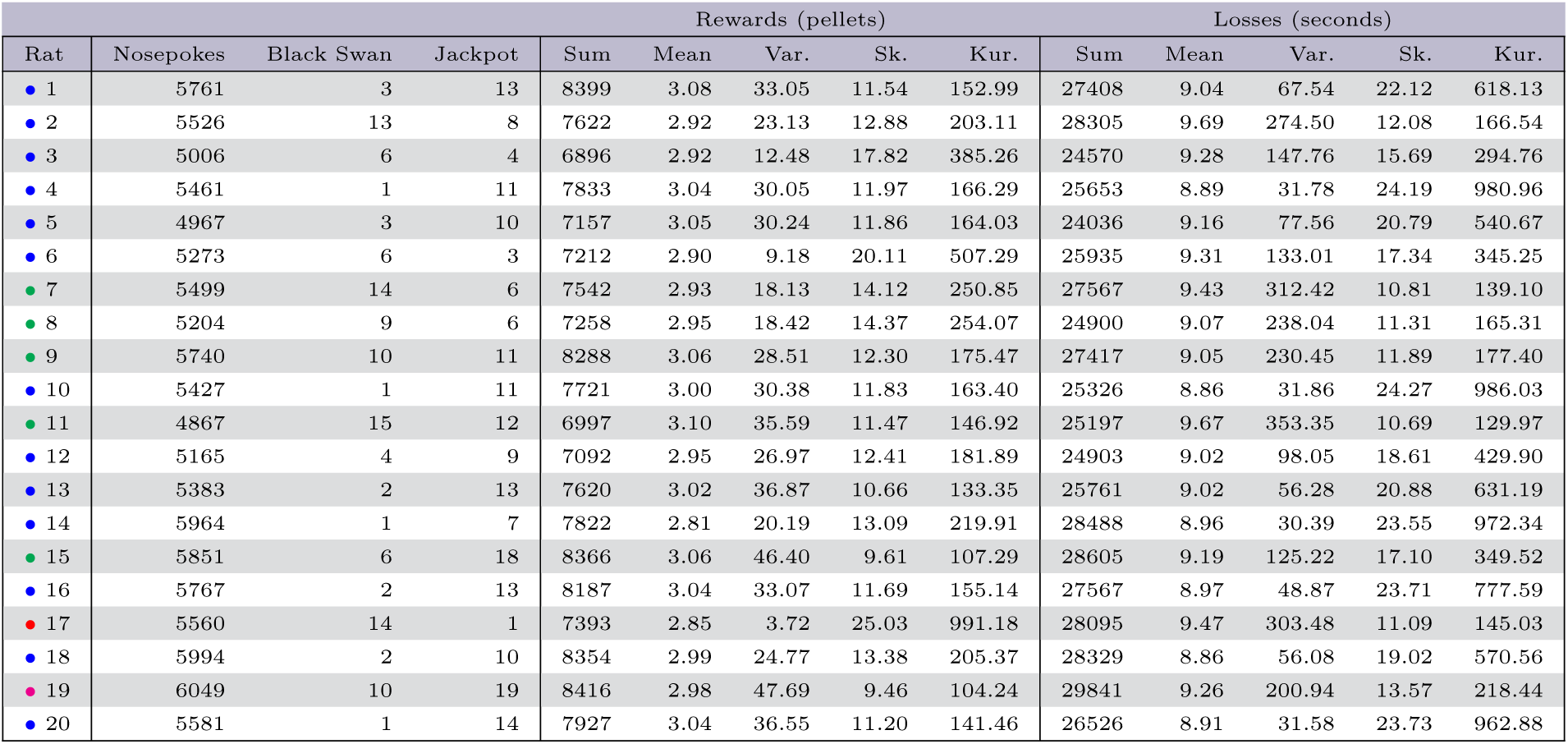
Ex-post total outcomes for each of the 20 rats over the final 41 sessions - summary statistics. Each row represents one rat and the color codes indicate the profile of the animal, as in Figure 2

### Convexity Premiums

Evidently the outcomes that we report in Table S11 depend on the convexity mix of options that each rat has chosen, as captured by our notions of Total and One-Sided Sensitivity. To go beyond Table S11 so as to capture the extent to which rats exploit convexity in their choices, it is perhaps informative to look at the outcomes for each rat in the following way. Right and left panels in Figure S1 refer to losses and rewards, respectively.

In each panel, rats are color-coded as in Figure 2, and they are represented horizontally by two segments which are normalized in the following way. Consider for example (blue) rat 1 in the first line. In the right panel about pellets, the point of the black segment most to the left represents the outcome that would have happened, had the rat chosen a concave option in the gain domain (that is, either Robust or Fragile), exclusively in all sessions. The point most to right corresponds, in contrast, to the counterfactual outcome in which all nose pokes of the rat correspond to a convex option (that is, either Vulnerable or Anti-Fragile). Superimposed on this background grey segment, is a bold-colored segment (blue because rat 1 is blue) that ends with a circle indicating what the rat has gained due to convexity in relative terms - what we call the *convexity premium* (see Appendix for more details). Rat 1 has a normalized convexity premium that corresponds to roughly 80% of what it would have got, in terms of sugar pellets, had it counterfactually chosen to mix only Vulnerable and Fragile, instead of Robust and Anti-fragile as it did. Similarly, one sees from the right panel of Figure S1 that the Anti-fragile rat 15 has a convexity premium that corresponds to roughly 90% of what it would have got, in terms of sugar pellets, had he chosen to mix only Vulnerable and Fragile. From panel (*b*) of Figure 2, we know that rat 15 has also, more rarely, picked the Robust and Fragile exposures and this is why its convexity premium in the gain domain is less than maximal. At the other extreme, the rat 6 has a convexity premium near zero. Note that two rats are outside the black segment in the gain domain, that is, have “negative” convexity premiums, and they are indicated by dashed lines. Rat 17 has picked the Anti-fragile exposure a few times but got only one extreme gain, while rat 14 is more sensitive and hence got more extreme gains but still too few of them. Therefore, both rats got less sugar pellets than what they would have gotten by mixing exposures that are concave in the gain domain (Robust and Fragile).

**Figure S1:**
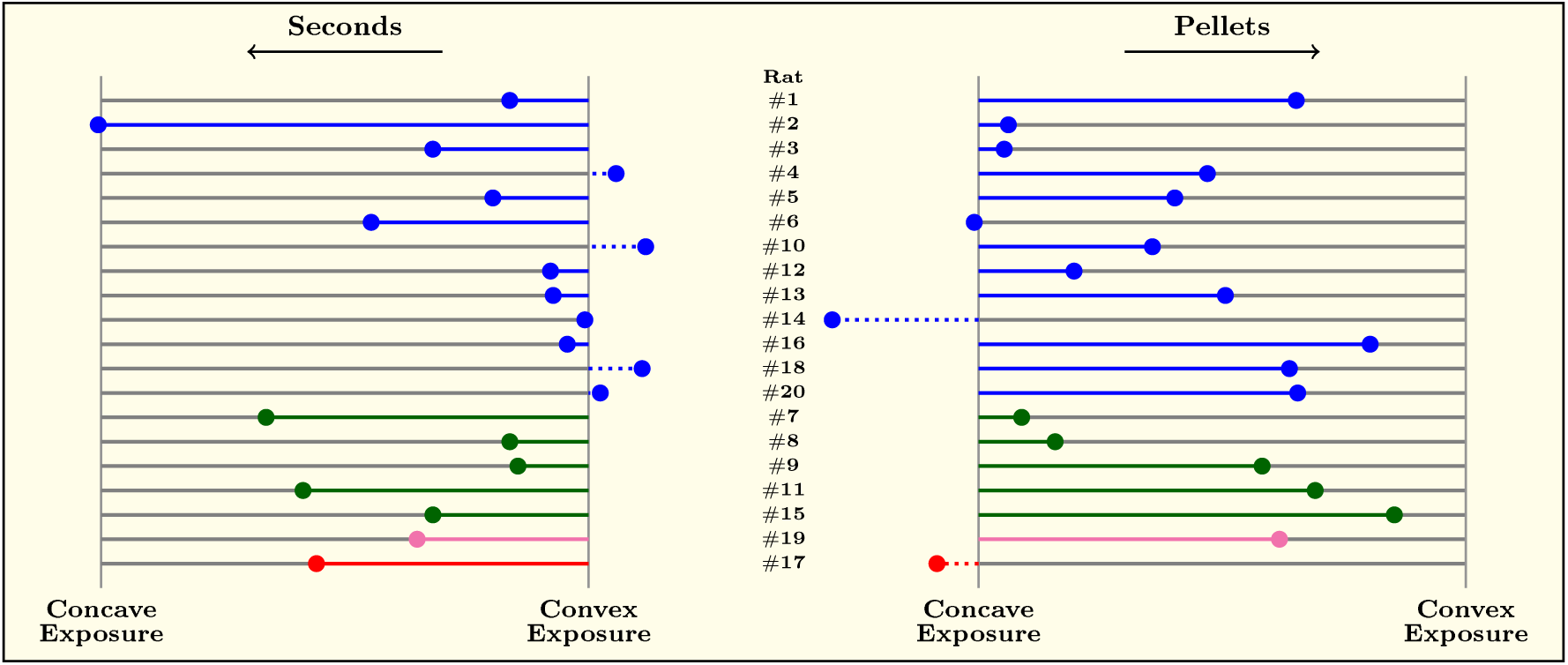
Convexity Premiums for each of the 20 rats (row) - see Appendix for details. Each row represents a rat and the color code indicates the profile of the animal, as in Figure 2. Towards the right, the dot materializes how many pellets were obtained relative to the number that would have been obtained if a convex menu had been chosen at each trial. Towards the left, the coloured dot indicates the total seconds of penalty obtained relative to what would have been obtained if a concave menu had been chosen at each trial.

The left panel in Figure S1 about time-out punishments can be read in a similar way, with now the point most to the left of the black segment corresponds to the largest loss in terms of waiting times, while the end point to the right corresponds to the lowest amount of time wasted. Therefore, the bold and colored segment indicates a negative premium: for instance, rat 2 on the first line in the left panel of Figure S1 has been exposed to the largest time-out punishment: it has a negative and large convexity premium because it mixed exposures with concave losses (that, is Fragile and Vulnerable), too often. Symmetrically, some rats turn out to get convexity premiums that are above 100% because they got too few black swans when they picked exposures that are concave in the loss domain). Comparing the right and left panels in Figure S1, one infers that *rats exploit convexity better in the loss domain than in the gain domain, that is, they more often avoid Black Swans than they get Jackpots*. This is of course consistent with our earlier observation that most rats exhibit moderate to high Black Swan Avoidance.

Dear Editors and Reviewers,

We thank you very much for giving us the opportunity to revise our submission. In the letter below, downloaded together with the article in the resubmission process, we reply to the reviewers.

We would like to express our surprise that the revised version of the manuscript was updated online as late as June 2026, whereas we submitted it in December 2025. In addition, we notice that we did not get feedback from Reviewer #1 about the revised manuscript, while Reviewer #2 essentially rephrased the same weaknesses, as if our point-by-point replies had not addressed them at all.

We would like to thank you again for allowing us on a path that we think has led to a much improved manuscript.

Best wishes,

Patrick Pintus, on behalf of all co-authors

Professor of Economics

Aix-Marseille University

**Reply to the Review #1**

Dear Reviewer #1,

Unless we are mistaken, we have not received your assessment of our point-by-point replies to all the weaknesses and recommendations, which were submitted together with the revised manuscript in the first step of the reviewing process.

It would be most useful to us, as well as to the general reader we presume, to have your feedback about whether we have satisfactorily addressed your main concerns and improved the revised version of the article that has been entirely rewritten.

**Point by point replies to the Review #2 (that appear in blue)**

Dear Reviewer #2,

We are not sure to understand the meaning of the first part of the review, which seems to be a repetition of the first report, with some variants and additions in the phrasing. Since we have replied in the first revision to all points raised then, simply copying and pasting here our replies, as they appeared in our first reply, seem redundant to us.

More generally, it seems unfortunate to us that the novelty of the design, approach (including some terminology) and results has received a broad negative assessment, despite our attempt to answer all your comments and recommendations in the first step.

Below appear our point by point replies to your review.

We thank you again for allowing us on a path that we think has led to a much improved manuscript.

**Public Reviews:**

**Reviewer #2 (Public review):**

**Summary:**

This paper attempts to examine how rare, extreme events impact decision-making in rats. The paper used an extensive behavioural study with rats to evaluate how the probability and magnitude of outcomes impact preference. The paper, however, provides limited evidence for the conclusions because the design did not allow for the isolation of the rare, extreme events in choice. There are many confounding factors, including the outcome variance and presence of less-rare, and less-extreme outcome in the same conditions.

**Strengths**

1. The major strength of the paper is the significant volume of behavioural data with a reasonable sample size of 20 rats.
2. The paper attempts to examine losses with rats (a notoriously tricky problem with non-human animals) by substituting time-outs as a proxy for losses. This allows for mixed gambles that have both gain and loss possible outcomes.
3. The paper integrates both a behavioural and a modelling approach to get at the factors that drive decision-making.
4. The paper takes seriously the question of what it means for an event to be rare, pushing to less frequent outcomes than usually used with non-human animals.

**Weaknesses**

1. The primary issue with this work is that the primary experimental manipulation fails to isolate the rare, extreme events in choice. As I understand the task, in all the conditions with a rare extreme event (e.g., 80 pellets with probability epsilon), there is also a less-rare, less-extreme event (e.g., 12 pellets with probability 5). In addition, the variance differs between the two conditions. So, any impact attributable to the rare, extreme event could be due to the less rare event or due difference in the variance (or other statistical moments, like skew or kurtosis). That the distributions can be shown to be different under specific assumption to value maximizing agents (e.g., with Jensen Gaps and Table 2) is not really relevant to what rats are sensitive and what drive their behaviour. The design here does not support the conclusions. Finally, by deliberately confounding rarity and extremity, the design does not allow for assessing the impact of either aspect on rat behaviour. Reply: This is a repetition almost word for word of point (1) appearing in your previous report, which we have addressed at length in our first-round reply. Therefore, we do not reiterate here to save space.
2. The RL modelling work also fails to show a specific impact of the rare extreme event. As best as I can understand Eq 2, the model provides a free parameter that adds a bonus to the value of either the two options with high-variance gains (A and V in the paper) or to the two options with high-variance losses (F and V in the paper). Or equivalently to the ones with “Jackpots” vs the ones with “Black Swans” (see Point 1 above as to how these different aspects are all confounded in this design). This parameter seems to only depends on whether this option could have possibly yielded the rare, extreme outcome (i.e., based on the generative probability) and was not connected to its actual appearance. [This point is unclear as the text says this, but the rebuttal states otherwise; plus some options never received the REE, see Table S11]. That makes it a free parameter that just bumps up (or down) the probability of selecting a pair of options. That may be due to presence of the REE or the other rare event or just the variance difference. Moreover, in the case of the “black swan” or high-variance loss conditions, this seems very much like a loss aversion parameter, but an additive one instead of a multiplicative one. Is there a theoretical claim here that “extreme losses” need an additive loss-aversion parameter? Reply: This is a repetition almost word for word of point (2) appearing in your previous report, which we have addressed at length in our first-round reply. Therefore, we do not reiterate here to save space.
3. The paper presented the methods and results with lots of neologisms and fairly obscure jargon (e.g., fragility, total REE sensitivity). That might it very hard to decipher exactly what was done and what was found. For example, on p. 4, the use of concave and convex was very hard to decipher; the text even has to repeat itself 3 times (i.e., “to repeat” and “in other words”) and is still not clear. It would be much clearer (and probably accurate) to say that the options varied along the variance dimension, separately for gains and losses. Option A was low-variance gains and losses. Option B was low-variance losses and high-variance gains. Option C was high-variance losses and low-variance gains, and Option D was high-variance losses gains. That tells much more clearly what the animals experienced without the reader having to master a set of new terminologies around fragility and robustness, which brings a set of theoretical assumption unnecessarily into the description of the experimental design. Alternatively, if the authors are wary of using the term “variance” because other moments of the distribution also differ, they could use “high-value gains” or “high-value losses” or something else which does not obscure the experimental design with jargon. Again, this goes back to point 1 above, whereby the different options differ on so many dimensions (as is made even more apparent in the rebuttal) that the design cannot isolate the impact of the variables of interest. Reply: This is a repetition almost word for word of point (3) appearing in your previous report, which we have addressed at length in our first-round reply. Therefore, we do not reiterate here to save space. The new words that we use are an attempt at reflecting the novelty of the approach and design.
4. Were the probabilities shuffled or truly random (seem to be fixed sequences, so neither)? What were the experienced probabilities? Given the fixed sequences, these experienced (“ex-post”) probabilities, could differ tremendously from the scheduled (“ex ante”) probabilities. It’s quite possible than an animal never experienced the rare, extreme event for a specific option. From Table S11, that is guaranteed to have happened in that 4 animals only ever experienced the “black swan” outcome once. It’s even possible (if they only picked a specific option on the 10th/60th choices by chance), that they only ever experienced that rare extreme event. This point still cannot be known given the information provided, which does not break down outcomes by options. The Supplemental in Table S11 only gives overall numbers but does not indicate what the rats experienced for each choice/option-which is what matters here. A simple table that indicates for each of the 4 options, how often they were selected, and how often the animals experienced each of the 6-8 possible outcome would make it much clearer how closely the experience matched the planned outcomes. In addition, by restricting the rare outcome to either the 10th or 60th activations in a session, these are not random. Did the animals learn this association? The text states that they did not, but no evidence is provided. Reply: This is a repetition almost word for word of point (4) appearing in your previous report, which we have addressed at length in our first-round reply. Therefore, we do not reiterate here to save space.
5. The choice data are generally presented in an overprocessed fashion with a sum and a difference (in both figures and tables). The basic datum (probability/frequency of selecting each of the 4 options) is not provided directly in the main text, even if it can theoretically be inferred from the sum and the difference. New right side of Table S4 is probably the most valuable piece in terms of explaining what rats did and should be highlighted a lot more. Inspection of that table reveals some interesting (and potentially worrying) results. Most notably, the vast majority of responding happens on the “anti-fragile” and “robust” option, often totalling around 90% of all selections, especially amongst the most common blue rats. Alas, those were all those the two options that were deliberately assigned to the two most preferred holes in the training phase (see p. 26). Does this reflect genuine preference for reward distributions or does this reflect a spatial hole bias? The assignment strategy makes this impossible to tell apart. Reply: This is a repetition almost word for word of point (5) appearing in your previous report, which we have addressed at length in our first-round reply. Therefore, we do not reiterate here to save space. In addition, in Figure 8 of Material and Methods, we report the changes in behavior between the training and final sessions. The related discussion stresses that half of the rats changed more than 30% of their choices in the final experimental sessions compared to the training sessions. See also our previous reply to point (8).
6. There is insufficient detail provided on the inferential statistical tests (e.g., no degrees of freedom or effect sizes), and only limited information on exactly what tests were run and how (bootstrapping, but little detail). Without code or data (only summary information is provided in the supplement), this is difficult to evaluate. In addition, the studies seem not to pre-registered in any way, leaving many research degrees of freedom. Not all studies need to be pre-registered and sometimes discovery of new things requires exploratory work, but preregistration does provide additional safeguards against overemphasizing post-hoc detected patterns-a serious issue in behavioural science. Moreover, this promotes transparency in reporting results and analyses, allowing for a better assessment of the strength of evidence for a claim. For example, here, were any alternative analysis pipelines attempted? Also, there were many sub-groupings of the animals and subsequent comparisons between them which all seemed post-hoc. On what grounds were these divisions made-were other divisions examined as well? Reply: This is a repetition almost word for word of point (6) appearing in your previous report, which we have addressed at length in our first-round reply. Therefore, we do not reiterate here to save space.
7. On p. 12 (Fig 4), there is an attempt to look at the impact of a rare, extreme event by plotting a measure of preference for the 10 trials before/after the rare, extreme event. In the human literature, the main impact of experiencing a rare, extreme event is what is known as the wavy recency effect (See Plonsky et al. 2015 in Psych Review for example, now cited). What this means is that there tends to there tends to be some immediate negative recency (e.g., avoiding a rare gain) followed by positive recency (e.g., chasing the rare gain). Typically, this refers to the specific option that yielded that outcome. First, as the other analyses do, the current analysis combines choice of the option that yielded the rare outcome with choice of other options, so that cannot directly assess the impact of the rare, extreme event on choice. Also, using a 10-trial window would thus obscure any impact of this rare, extreme event. There is mention of the very next trial, but an analysis that looks at the 10-trial time course trial-by-trial could reveal any impact that might be predicted from the human literature. Reply: This is a repetition almost word for word of point (7) appearing in your previous report, which we have addressed at length in our first-round reply. Therefore, we do not reiterate here to save space.
8. As I understood the method (p. 31), the assignment of options to physical locations was not random or counterbalanced, but deliberately biased to have one of the options in the preferred location. This would seem to create a bias towards a particular option and a bias away from the other options, which confounds the preference data in subsequent analyses. Table S4 reinforces this concern where the vast majority of response are clustered in the two most preferred options from training. Reply: This is a repetition almost word for word of point (8) appearing in your previous report, which we have addressed at length in our first-round reply. Therefore, we do not reiterate here to save space.
9. Are delays really losses? This is a big assumption. Magnitude and delay are different aspects of experience, which are not necessarily commensurable and can be manipulated independently. And, for the model, how were these delays transformed into outcomes for the model. Eq 1 skips over that. Is there an assumption of linearity? In addition, I was not wholly clear if the delays meant fewer trials in a session or if the delays merely extended the session and meant longer delays until the next choice period. Reply: This is an exact repetition of point (9) appearing in your previous report, which we have addressed at length in our first-round reply. Therefore, we do not reiterate here to save space.

**Other points**

1. I think the authors still misunderstand the concept of “hot-stove effects”. The idea is that the experience of a very bad outcome can lead to avoiding the situation again (i.e., not sampling that option) and can provide the appearance of oversensitivity to that bad outcome. Here, that might be more thought as “black-swan avoidance”. Imagine if, to the rat, all options are equal in value, then some initial bad luck in encountering the black swan might make the animal avoid that option, even though with enough experience, then it would have been equal in value. Reply: This point is reminiscent of recommendation (5) appearing in your previous report, which we have addressed at length in our first-round reply. Therefore, we do not reiterate here to save space.
2. I am still not convinced that the Jensen inequalities add to this paper in terms of understanding the rat behaviour. That may be more suited for a different paper about the statistical and mathematical properties of certain generative distributions, but not here given what rats actually choose and experience. Reply: This point is reminiscent of recommendation (8) appearing in your previous report, which we have addressed at length in our first-round reply. Therefore, we do not reiterate here to save space.
3. Providing the data open access is very good. The code, however, should be equally available and not just upon request. Code needs to be available for assessment during peer review and for reproducibility checks. There are substantial enough problems with reproducibility in the field that code availability should be a minimum criterion for publication (see Miske et al., 2026 in Nature for the most recent large-scale evaluation of this problem). Reply: This point is reminiscent of recommendation (9) appearing in your previous report, which we have addressed at length in our first-round reply. Therefore, we do not reiterate here to save space.
4. The paper still somewhat mischaracterizes the literature on rare events, posing it as a series of “exceptions”, rather than recognizing that a huge chunk of the literature uses rare events rarer than 10%. Also, there is even existing terminology in that literature for exactly the situation that is being created here-rare treasures (aka jackpots here) and rare disasters (aka Black Swans here). Reply: This is reminiscent of point (10) appearing in your previous report, which we have addressed at length in our first-round reply. Therefore, we do not reiterate here to save space.
5. Defining the observed behaviour in terms convexity, instead of stating choices more plainly obscures what is done/found. This is especially the case here because convex and concave mean different things when applied to gains/losses in terms of whether or not that option can lead to the REE. The use of the terms obscures rather than clarifies and probably is best left for the discussion (and maybe the intro) when mapping from theoretical distributions to the experiment at hand. In the paper, even the bottom of p.5 seems to incorrectly define “Total Sensitivity” as the combined proportion of selecting convex options in either domain, which does not map how convex is defined in Fig 1B or elsewhere in the text. Reply: This is reminiscent of point (3) appearing in your previous report, which as stated above we have addressed at length in our first-round reply. Therefore, we do not reiterate here to save space.
6. Fig 1C is baffling. Why are probabilities drawn moving away from the origin? The standard scientific plotting convention is for numbers to grow when moving away from the origin. That would be vastly clearer. Also, the color coding is confusing. Green-red maps onto convex-concave, but that would naturally seem to indicate gains vs losses, not convex vs concave. And why are probabilities growing larger in both directions from the origin? Much more sensible to communicate the procedure would likely be a standard plot of magnitude vs probability. Reply: As explained in the main text of the article, panel (c) in Figure 1 reveals an important and novel feature of the design: gains and losses are either convex or concave in **decreasing** probabilities, meaning that the gain/loss gets more rare, it becomes larger (if convex) or smaller (if concave) in a nonlinear fashion.
7. Discussion: I think the main difference between the human situations discussed and this experiment is that humans have not experienced those rare “black swan” outcomes. Rather, they hear about the disasters that are possible and do not incorporate that information, as discussed in the description-experience literature already cited in this paper (though not in that context). Reply: We thank the referee for pointing this out. However, it is not clear to us that the various trauma experienced in real life by humans do not share properties with Black Swans in our experimental design.

**Recommendations for the authors:**

**Reviewer #2 (Recommendations for the authors):**

**Small points:**

- p. 4. Options are not strategies. Picking two out of the 4 options more often, is not picking two strategies. That’s picking two options (also on p. 7, but really throughout).

Reply: This has been changed in the text.

- The sums don’t add up in probability of outcomes. How were they 0.9, 0.1, 0.01? What were they exactly? What exactly is epsilon?

Reply: Exact probabilities, which of course add up to one, are given in Material and Methods (page 29).

- The term “one-sided sensitivity” to refer to the selection of options include rare-extreme outcomes is also confusing. There are many other words that might work better here, including “encounter”, “choice”, “selection”, or even “preference”.

Reply: One-sided Sensitivity if a proposed measure of the choice asymmetry, as explained in the text (pages 5-6, including caption of Figure 1).

- I don’t think that Figure 3 is more intuitive. It’s still a summed measure, with rats sub-divided into post-hoc groups. Also, aren’t the groups divided precisely in part by the selections that are being shown in this plot?

Reply: There is no claim that Figure 3 is more intuitive. As explained in Material and Methods (pages 33-34), it provides an alternative way to represent behaviors (BSA/JPS vs TSREE/OSREE).

- Fig 3 has no black or dark and light grey. There is no one asterisk in the figure. The light grey is irrelevant because all bars must sum to 1. No error bars or equivalent on the plot. Reply: Dark and grey bars sum to one but have a different interpretation, as explained in the text (pages 10-11). We have dropped the one asterisk reference.

- p. 14. Technicality, but alpha need not be bound at 1 for a convergent equation. It only needs to be bound at 2 (real maximum value). Above 1 (but below 2) would be a type of overcompensation that would also converge at same Q value.

Reply: The assumption that alpha is positive and bounded by one is common in the literature and is important for stochastic convergence. Assuming that alpha is between one and two has undesirable implications, beyond possible non-convergence, in particular that Q values may fluctuate.

- p. 24. What was the generative frequency (not experienced frequency). Not adequately explained. Less than half a percent of all outcomes? Of all times that the relevant choice was made. This is not explained.

Reply: The sequences used to generate gains and losse are detailed in the Supplementary Material.

- p. 4. Says 41 sessions, but methods say 40 (e.g., Fig 7 and surrounding text)

Reply: This has been corrected in the text.

- My understanding of Tables S5 and S6 is that the best models did not include REE in Q sub-values. That seems to be opposite to the textual conclusion.

Reply: As stated in Table 1 (page 15), which summarizes results in Tables S5-S6, the selected models for rats 3, 11, and 12 do not integrate REE while the selected models for rats 4, 16 and 20 integrate REE though Q values and REE decision weights. For most rats, REE are integrated in the selected models through the latter only.

- What are the parameters in Table S7? The full 8-parameter model? But how are these the ones selected by BIC as per caption? Also, there are a number of anomalous values-lots of 1s and 0s, plus a +40 on the value of the losses for Rat 20. Columns are always gain then loss except first column.

Reply: we report in Table S7 all estimated parameters for each rat, but only for the model selected by the BIC criterion (see Table S5). Admissible values for alphas include 0 and 1. All estimates have been checked for identification. The first and second columns have been switched to have gain first.

- Table S11 does not sub-divide by option selected. So, we cannot tell if the rats actually experienced the outcome distribution associated with some options.

Reply: Given the large number of nosepokes generated by each rat, reporting all actual gains and losses would consume too much space. On the other hand, given that each option has only three frequent and rare values, it is expected that each will be experiences by each rat, in view of the choice frequencies reported in Table S2 (page 52).

## Notes

https://github.com/PatrickPintus/Rare-and-Extreme-Events

### Competing Interest Statement

The authors have declared no competing interest.

### Summary of Updates

we here have added the replies to the reviewer's comments after the first revision. We also attached the point per point replies showing how redundant they are with the first round of review.

